# Conserved upper thermal limits and small safety margins in soil copiotrophic bacteria

**DOI:** 10.64898/2026.02.06.703311

**Authors:** Ariel Ignacio Favier, Alejandra Hernández-Terán, Celia C. Symons, María Rebolleda-Gómez

## Abstract

One of the key uncertainties in climate change models is how microbes will adapt to rising temperatures. Large-scale comparisons of bacterial thermal performances show a clear boundary between mesophiles and thermophiles. Here, we investigated whether phylogenetic constraints limit the adaptive potential of bacteria to warming soils. Focusing on copiotrophs within Gammaproteobacteria, we found that both thermal optima and upper thermal limits are constrained; variation in these traits decreases above 42°C across the phylogeny, with minimal influence of present-day bioclimatic variables. This, along with the reduced thermal safety margins found in fluctuating hot climates, suggests that many isolates may already be maladapted to local temperature variability. Our findings indicate that these constraints interact with the geometry of thermal performance curves, imposing a trade-off between high-temperature performance and the risk of substantial fitness losses above Topt. Overall, this work underscores potential limits to thermal adaptation and their implications for bacterial fitness.

## Introduction

The adaptability of organisms to a warming planet is one of the most pressing questions in ecology, as average temperatures and thermal variability increase at unprecedented rates (Zeebe and Zachos 2013; Zeebe et al. 2016). Soil microbial communities, key in regulating terrestrial carbon fluxes (Trivedi et al. 2013), are expected to increase their respiratory activity in warming soils, creating a positive feedback loop with warming (Cox et al. 2000). However, this prediction implies an unconstrained upper limit to microbial activity across taxa and environments (Alster et al. 2016, 2020), which contrasts with the evidence of strong limits on mesophilic adaptation attributed to the instability of cellular structure and physiology at supra-optimal temperatures (Corkrey et al. 2016).

Bacteria readily evolve higher thermal tolerance as a result of experimental evolution in the laboratory (Tenaillon et al. 2012; White et al. 2025; Toll-Riera et al. 2022; O’Donnell et al. 2018; Bennett et al. 1990). Nevertheless, when examining bacterial growth rates across different temperatures, there appears to be a clear gap between mesophiles and thermophiles between 45°C and 50°C (Corkrey et al. 2016; Bausum and Matney 1965). The physiological and evolutionary constraints of these limits remain poorly understood. Similarly, long-term experimental warming studies (Frey et al. 2013; Melillo et al. 2017) show variable trends of metabolic efficiency in response to warming. In these experiments, it is often impossible to separate the effects of changes in resources, microbial community composition, and physiological and evolutionary responses of individual species. A more systematic understanding of microbial thermal physiology and its evolvability is important to start untangling these processes.

At the organism level, thermal physiology can be described by a thermal performance curve (TPC), which models functional metrics in relation to temperature. For ectotherms, the shape of the TPC is characterized by an increase in performance from a lower critical temperature (CTmin) to a thermal optimum (Topt), where organisms achieve peak performance (AUCmax); usually followed by a steeper decrease until reaching the upper critical temperature (CTmax), above which mortality exceeds reproduction (Angilletta et al. 2010) (Fig. 1A). TPCs are concave down at high temperatures and thus, average performance across temperatures is lower than performance at the average temperature (Jensen’s inequality). That is, if an organism evolves a thermal optimum that tracks the average temperature of a location, its average performance over the temperature variation of the environment will be lower than if it tracked this variation. The opposite pattern holds for low temperatures (where the TPC is concave up). Because of this geometry, it is expected that, in variable environments, organisms will evolve higher thermal optima than the average environmental temperature (Amarasekare and Johnson 2017; Bernhardt et al. 2018; Martin and Huey 2008).

**Figure 1:**
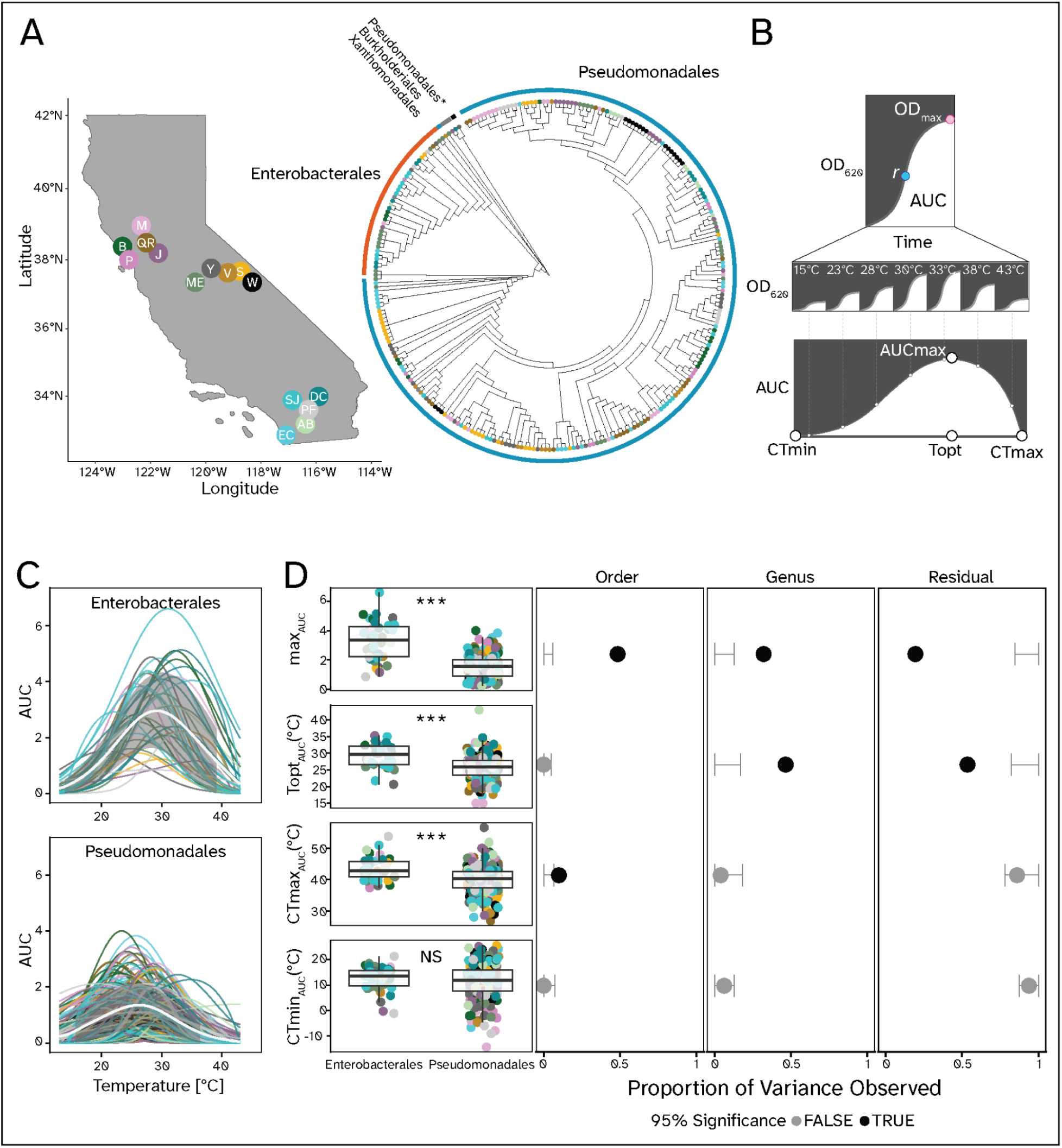
Thermal traits are differentially conserved in a collection of bacterial isolates from Californian topsoils. **(A)** We collected topsoil samples from fifteen sites and isolated bacteria at three temperatures (left). We sequenced a random sample of four hundred isolates and constructed a phylogeny (center). Points are colored to represent the sampling site, and the ring around the dendrogram is colored according to Order. The “*” signals a representative of the family Moraxellaceae, which consistently grouped outside of the main branch of Pseudomonadales. **(B-C)** We obtained growth curves after growing each isolate in M9 media supplemented with D-Glucose for 24 hs, and we then fitted TPCs using the Area Under the Curve (AUC) for the two majoritary Orders (right). Shaded trendlines signal the mean SD of AUC across temperatures. **(D)** We compared the thermal traits between Enterobacterales and Pseudomonadales (left) using one-way ANOVAs (*** = p < 0.001, ** = p < 0.01, * = p < 0.05, NS = p >= 0.05), and we then compared the variance observed at each relevant taxonomic level to a confidence interval (CI) of variance expected after permuting trait values across the collection (nsim = 10000) (right). Black points outside the CI for the Order or Genus columns represent phylogenetic signal at that taxonomic level, while grey points represent the opposite. Site abbreviations are as follows: W - White Mountain Research Center, SJ - James San Jacinto Mountains Reserve, S - Sierra Nevada Aquatic Research Laboratory; V - Valentine Camp, Y - Yosemite Field Station, P - Point Reyes Field Station, B - Bodega Bay Research Center; AB - Steele/Burnand Anza-Borrego Desert Research Center Research Center, DC - Boyd Deep Canyon Desert Research Center, EC - Elliot Chaparral Reserve, QR - Quail Ridge Reserve, PF - Pinyon Flats; ME - Merced Vernal Pools and Grasslands Reserve, M - James McLaughlin Natural Reserve, J - Jepson Prairie Reserve.

The extent to which the TPCs of ectotherms can adapt to increases in temperature and temperature variability will depend on the existing diversity and evolutionary lability of their different thermal traits. While novel mutations can modify thermal responses of organisms ( (Toll-Riera et al. 2022; Rodríguez-Verdugo et al. 2014), associations between different thermal traits and conserved physiological responses can limit adaptation to increased temperatures.

Genetic variation within populations is correlated to species divergence and linked to the ability of populations to respond to rapid environmental change (Holstad et al. 2024). Low phylogenetic variance across environments is often an indicator of evolutionary constraints (McKitrick 1993).

Soils harbor an almost experimentally intractable bacterial diversity, dominated by mostly yet uncultured (Lok 2015) members of the phyla Proteobacteria, Acidobacteria, and Actinobacteria (Janssen 2006; Delgado-Baquerizo et al. 2018). Within Proteobacteria, the class Gammaproteobacteria covers a great taxonomic and functional diversity, including taxa highly responsive to temperature fluctuations in soils (Tikhonova et al. 2019; Tao et al. 2020; Goldfarb et al. 2011; Williams et al. 2010). Members of this class can be classified as copiotrophs (Finn et al. 2021; Fierer et al. 2007; Koch 2001), often readily culturable in standard laboratory settings due to their high nutrient acquisition ability (Stone et al. 2023; Malik et al. 2020), making their physiological measurements practical. Moreover, copiotrophs are more likely to be limited at higher temperatures, as such conditions favor slower-growing, more stress-resistant microbes (Abreu et al. 2023; Du et al. 2024).

To test the influence of phylogeny on the thermal traits of copiotrophic bacteria, we measured thermal performance of a collection of isolates within Gammaproteobacteria from topsoils across fifteen sites in California spanning a broad range of climate conditions. We tested 1) the level of conservation of thermal traits, 2) their correlation with environmental factors, and 3) the potential for phylogenetic constraints to limit adaptation to increased warming and environmental variability. We hypothesized that performance peak values at the Family and Order levels would be phylogenetically conserved, reflecting different substrate preferences. In contrast, we expected thermal traits, such as the thermal optima and critical temperatures (Malusare et al. 2023), to show more variability and positively correlate with site mean annual temperatures (Thomas et al. 2016) but have a larger safety margin (*i.e.,* difference between the thermal optima and the average temperatures) in variable environments. Overall, we found striking conservation of thermal optima and performance peaks, clear differences in TPC shape between taxa, and a strong upper critical temperature limit near 42°C across clades, which suggests a limited adaptive potential for Gammaproteobacteria in warmer soils.

## Materials and Methods

### Sampling and Isolation

We sampled surface soils (∼50 mm) from 15 sites spanning a wide range of climates across California, from Spring to Fall of 2022 (Fig. 1A). We collected five samples from each site, trying to capture the variation in solar exposure and associated vegetation. We then incubated 5 g of each sample in PBS with cycloheximide for 24 hs at three different *temperatures of isolation* (15°C, 25°C, and 35°C). Afterward, we plated the samples on chromogenic agar (Universal Differential Chromoselect Agar, Millipore Sigma; casein 4 g/L, peptone 15 g/L, and a proprietary chromogenic mix) and incubated them at their respective *temperature of isolation*. We randomly selected 686 potential bacterial isolates, and sequenced the whole genomes of 400 at SeqCoast Genomics (Portsmouth, NH, USA). These isolation protocols are inherently biased toward obtaining representatives of copiotrophic taxa. We used the genomic information to build the phylogenetic tree (Supplementary Methods).

### Environmental data

We gathered environmental data from weather stations at all the sampled sites. For all sites except the Merced Vernal Pools and Grassland Reserve (MVPGR) and Pinyon Flats/Pinyon Crest (PF), we obtained climate data from the Dendra database (https://dendra.science/). We obtained MVPGR data from the San Joaquin River weather station (https://water.ca.gov/) and PF from the Boyd Deep Canyon weather station at Pinyon Crest (https://deepcanyon.ucnrs.org/weather-data/; only monthly temperature minima and maxima, and monthly average rainfall were available for this station).

From each station, we obtained the daily 50mm - 100mm soil temperature minima, maxima, and average corresponding to the time series 2002-2022 (when available). We calculated the temperature soil bioclimatic variables (SBIO1 - 7) as implemented in previous studies (Supplementary Table 1)(Lembrechts et al. 2022). The bioclimatic variables we obtained often co-varied. To reduce dimensionality and summarize covarying climate variables, we used a principal component analysis, grouping our sampling sites in broader climate categories.

Precipitation data from the UCNRS database were only cumulative and not present for all sampling locations. Therefore, we obtained monthly precipitation data of all the sampling sites from the <u>PRISM</u> f. After comparing the PRISM data with the available UCNRS results, we found no significant differences (Fig. S1), and thus we used only PRISM precipitation data for all subsequent analyses.

Finally, we calculated the plant growth season per site as the time interval around the annual maximum positive rate of change NDVI (Normalized Difference Vegetation Index). While a positive NDVI signals the presence of green vegetation against a non-vegetated background, a positive derivative indicates increases in greenness, or plant growth. We downloaded 500 m point sample NDVI measurements at a 16-day resolution for all our sampling sites from the NASA AppEARS (https://appeears.earthdatacloud.nasa.gov) website for the period 2013-2022.

### Culturing methods

We measured the growth curves of our isolates at a minimum of six temperatures covering the mesophilic temperature range (∼ 13 - 45°C). In most studies, it is hard to untangle the effects of temperature and of changes in carbon preferences and availability. Therefore, we decided to use defined media (M9) with a single carbon source (D-Glucose at 0.02%) for all of our growth assays. Glucose (Glu) is a simple carbon source that can sustain the growth of most culturable heterotrophic Gammaproteobacteria. We ran additional tests on a subset of isolates to evaluate the impact of a 10-fold increase in glucose concentration on thermal performance and found only changes in performance peak, but not horizontal shifts in thermal limits or optima (Fig. S2). Briefly, we grew the isolates in 96-well microplates with 400μL of rich media (TSB) for 24 hs at 25°C, before transferring 20μL of them to a preculture of 180μL of M9+Glu media at the target temperature (+-2°C) for acclimation. After 24 hs, we transferred 4μL of each preculture to a plate with 196μL of fresh M9 at the target temperature and measured the growth kinetics of the plate for 24 hs without agitation. We obtained up to 3 experimental replicates per temperature.

### Growth curve and TPC fitting

We used the R package *gcplyr* (Blazanin 2023) to fit models to our growth curves and obtain their main growth parameters. After removing outliers and contaminants (Supplementary Methods, Fig. S3 and S4), we used the R package rTPC (Padfield et al. 2021) to fit TPCs for isolates with growth curves at six or more temperatures (moderate to high resolution) (Kontopoulos et al. 2024). We used five phenomenological models (briere2_1999, gaussian_1987, rezende_2019, spain_1982, lactin2_1995) that often outperform mechanistic TPC models when a moderate or high number of experimental temperatures is measured; these models also have few parameters, which makes fitting easier as well (Kontopoulos et al. 2024). Since no individual model performed best across the collection, we then used a weighted AICc approach to average fits. We calculated the TPCs for growth rate (*r*) and the area under the fitted curve (AUC) for each isolate. Despite the overall similarity in TPC shape across indicators, *r* was more variable across replicates and sensitive to differences in fit. AUC provided a more stable estimate of growth, while also capturing the total yield (ODmax). Thus, we used AUC-derived TPCs for further analyses. After filtering out unrealistic fits (see Supplementary Methods, Fig. S5), we retained 244 TPCs for further analysis.

### Statistical analyses

We ran all statistical analyses in Rstudio (v.2023.12.1.402), running R version 4.3.2. To estimate the separate effects of phylogeny and environmental factors on each thermal trait, we fitted a linear mixed regression model of the form *lmer(Trait ∽ PC1 + PC2 temperature_of_isolation + (1|Order/Genus))* using the package *lmerTest* (Kuznetsova et al. 2017). In this model, we first calculated the marginal effects of the components of environmental variation across our sites and the temperature at which we isolated different strains (temperature_of_isolation) on a given trait. We extracted partial R2 values for each fixed effect using the package *partR2* (Stoffel et al. 2021), and p-values using the Satterthwaite approximation. Afterwards, we calculated how much of the residual trait variance could be explained by the effects of two nested taxonomic groupings following methods designed to estimate phylogenetic signal (Lennon et al. 2012). We excluded Family from the analysis because its distribution was nearly identical to that of Order. We also removed Species because many isolates lacked a taxonomic classification at that level. We then simulated a null distribution of trait variance (trait values are randomly distributed across the phylogeny) by recalculating the observed variance after permuting the trait values within our collection (Number of Permutations = 10000). We considered a trait to show significant phylogenetic signal when the variance at a given taxonomic level fell outside the 95% confidence interval of the null distribution.

For comparing trait means between specific clades, we used the function *aov*, followed by post-hoc comparisons (functions *TukeyHSD* and *emmeans*, respectively), with a significance level of 95%. We also compared the dispersion between linear regressions (function *lm*) using F-tests, and the slopes using Welch’s t-tests. We estimated pairwise correlations between variables using the *cor* function from the *stats* package. We generated all plots using the *ggplot2* package and compatible additional packages, such as *corrplot* (Wei et al. 2017) and *ggtree (Yu et al. 2017)*.

## Results

### Thermal optima and performance peaks are conserved across taxa

We aimed to determine the relative contribution of phylogenetic similarity and local climate in shaping thermal performance across copiotrophic Gammaproteobacteria from different soils across California. Most of our isolates belonged to Orders Pseudomonadales and Enterobacterales, and were widely distributed across our sampling sites (Fig. 1A). We hypothesized that strong phylogenetic constraints would lead to large differences in thermal traits across taxa, regardless of the local environment. Consistent with this hypothesis, we saw significant differences in TPC shape reflecting phylogenetic relatedness (Fig. 1C). After accounting for the marginal effects of environmental variation components and isolation temperatures, performance peak showed significant phylogenetic signal at the Order level compared to random expectations (Variance observed = 48.5%; Fig. 1D). We observed significantly higher values of performance peak (p < 0.0001, Cohen’s d = 1.98), and CTmax (p-value = 1.05e-05, Cohen’s d = 0.78) in Enterobacterales compared to Pseudomonadales (Fig. 1C-D).

Topt values were significantly conserved at the Genus level (Variance observed = 46.5%; Fig. 1D), a trend driven by the genera Enterobacter and Mixta, which consistently displayed higher values (Fig. S6). Although the phylogenetic signal was higher at the Genus level, the Order Enterobacterales displayed significantly higher thermal optima than Pseudomonadales (p_value < 0.0001, Cohen’s d = 1.01; Fig. 1D). Despite significant phylogenetic conservation, residual differences still explained a significant proportion of the variation in Topt (38%). In contrast, phylogenetic differences did not explain the variance in thermal limits beyond what would be expected under a randomized trait distribution. CTmax showed low variance, partially explained by conservation at the Order level (10%), while CTmin was highly variable with no detectable phylogenetic signal.

### Experimental isolation temperature and environmental variables explain a modest fraction of thermal trait variance

A lack of phylogenetic conservatism could be due to environmental adaptation overriding phylogenetic differences. We evaluated the contribution of the two principal components of environmental variation on each of our thermal traits of interest. We first characterized each site according to bioclimatic variables (see Methods). Two axes (PC1 and PC2) explained 61% and 27% of the variation in bioclimatic variables across sites; these axes were broadly associated with temperature variability and average, respectively (Fig. 2A). Due to their strong negative correlation with temperature means and temperature variability, precipitation variables did not significantly change the point distribution along these two axes, and thus we removed them from further analyses (Fig. S7). Based on the values along these two main components of variation we separated our sites into four general categories: hot variable sites (Quail Ridge and desert sites), cold variable (high altitude sites in the sierras and Mt. San Jacinto in Southern California), hot stable (inland northern sites), and cold stable (coastal northern and lower altitude sites in the sierras).

**Figure 2:**
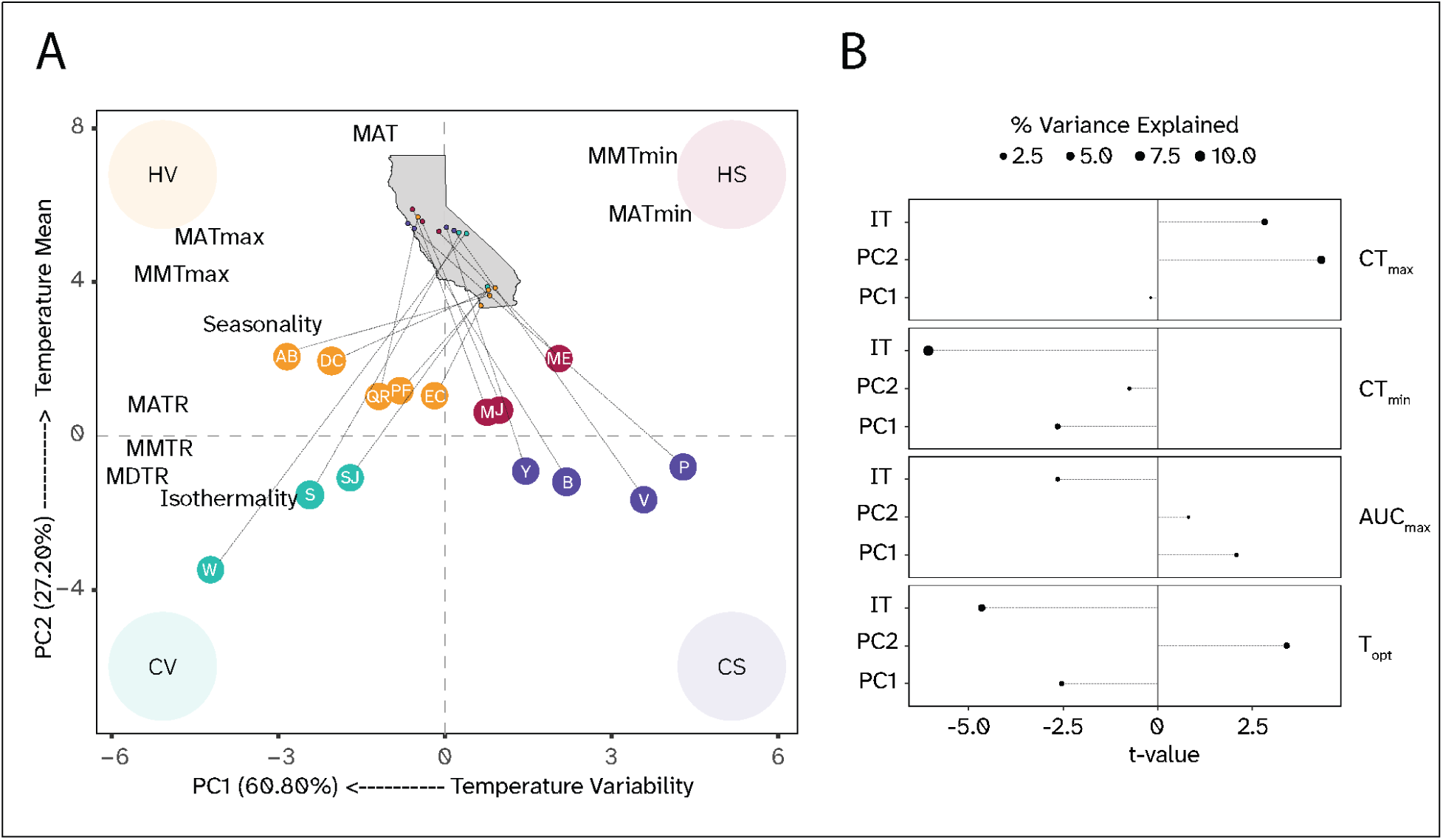
Principal Components of environmental variation explain a small amount of thermal trait variance. **(A)** PCA of soil bioclimatic variables across sampling sites. PC1 had a high negative correlation with metrics of temperature variability across temporal scales. PC2 was positively correlated with increases in mean annual temperatures, as well as minima and maxima. This allowed us to categorize our sites based on their position into four categories: Cold Variable (CS), Cold Stable (CS), Hot Variable (HV) and Hot Stable (HS) (site abbreviations are as in Figure 1). **(B)** The t-value denotes the magnitude and sign of the association between a variable and a thermal trait, while the point size represents the trait variance explained by such variable. Overall, we see that experimental conditions explained a consistently higher fraction of trait variance than broad environmental variation components, and that the sign of the relationship depended on the variable and trait.

We expected Topt and CTmax to be associated with mean and maximum temperatures, and CTmin with the minimum temperatures, while increased variability would lead to a broader thermal niche and higher Topt and CTmax to increase the safety margin and maximize performance across time. Instead, PC1 and PC2 only explained a small proportion of the variance in thermal traits across our collection (Fig. 2B; Supplementary Table 2). As expected by its phylogenetic conservatism, the performance peak was not explained by isolation temperature or environmental components (Combined explained variance = 1%). Approximately 5% and 2% of the variance in CTmax and Topt could be explained by PC2, associated with increases in mean temperatures, whereas PC1, negatively associated with temperature variability, explained a negligible 1.6% variance in CTmin. The experimental isolation temperature was negatively correlated with CTmin and Topt, and explained a higher fraction of their variance (10% and 4%, respectively) than PC1 and PC2 combined. That is, we observed a slight upward shift in thermal traits with mean temperatures, but temperature variability did not significantly influence thermal traits.

### Limited variation in thermal maximum suggests constraints in adaptation to high temperatures

To better understand potential constraints shaping the evolution of different thermal traits, we first compared variability across isolates for each thermal trait. CTmin and the performance peak were overall more variable than Topt and CTmax (Fig. 3A). This trend persists within each climate profile (Fig. S8) and indicates a potential global constraint to the evolution of Topt and CTmax.

**Figure 3:**
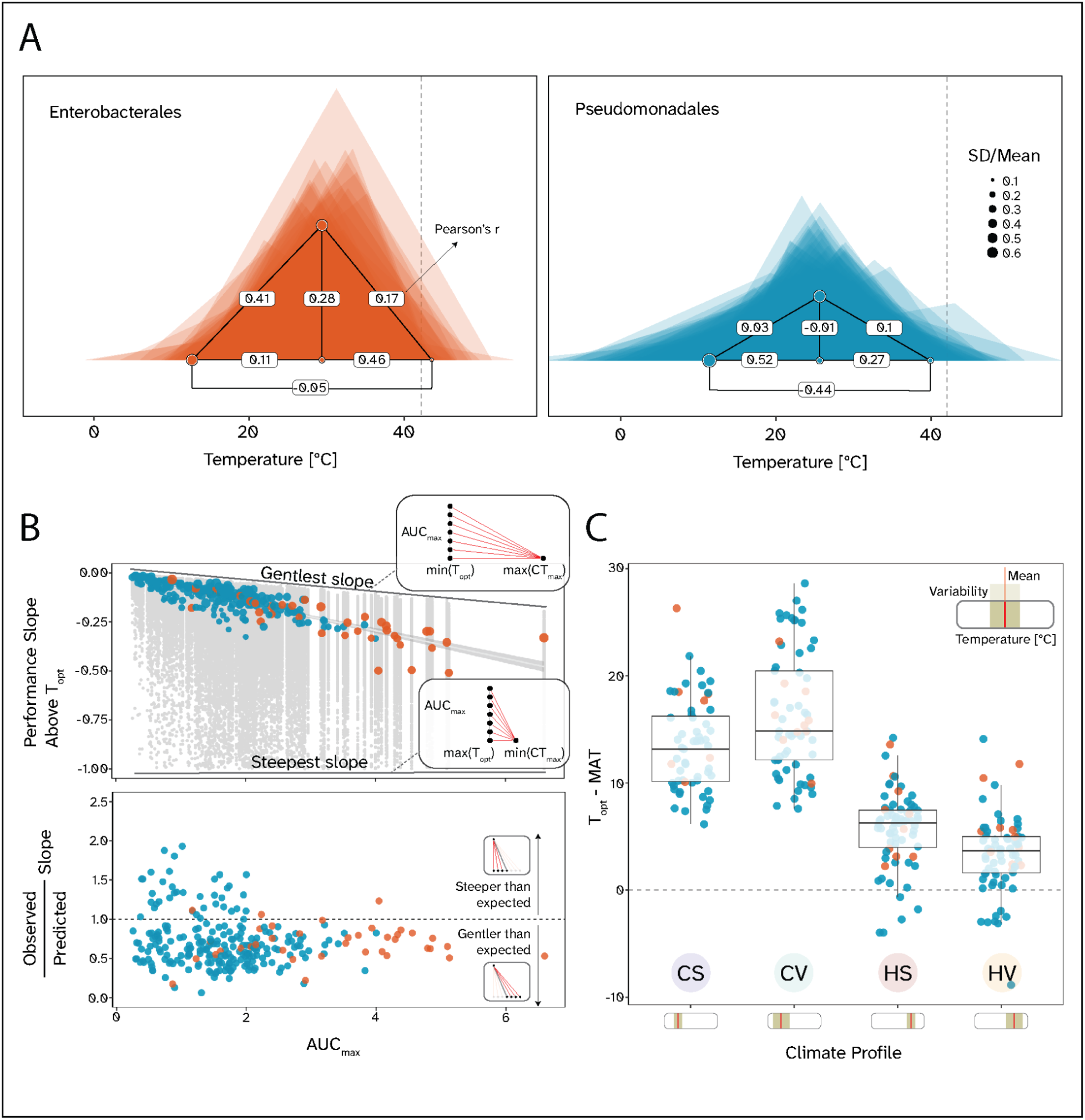
Variation in TPC shape is limited to changes in the lower thermal limit and performance peak. **(A)** We generated simplified TPCs for Enterobacterales (Orange) and Pseudomonadales (Blue), connecting the main traits represented by points (CTmin, CTmax, Topt, and Peak) with straight lines to form a triangle for each curve. The dashed line indicates the upper thermal limit documented for mesophilic enzymes at 42°C. Translucent triangles represent individual isolates, and full triangles represent average values by taxon. Point size is proportional to the trait coefficient of variation (CV) within that taxon. The labels on the lines connecting traits represent the correlation (Pearson’s r) coefficient between them. **(B)** We then calculated the slope of performance decrease above the thermal optimum of each simplified TPC. Here, we show that the observed slopes (colored points) become steeper with performance peaks, suggesting the apparent robustness of the upper thermal limit. Compared to slopes obtained from a randomized distribution of thermal traits keeping the original peak values (grey points and dashed trendline), the trend of observed slopes (red dashed line) shows a higher goodness of fit than any set of simulated slopes, and lands closer to the line of gentlest possible slopes, as represented by the ratio between the observed and predicted slopes at each value of performance peak (bottom). **(C)** Finally, we evaluated the hypothesis of Jensen’s inequality-based increases of Topt with temperature variability in cold and hot climate profiles. We observe contrasting trends: the difference between Topt and mean annual temperatures (MAT) increases with variability in generally cold environments, while it decreases in hot environments.

To avoid fitting biases, we compared variation in growth across temperatures, and observed a decline in overall growth and growth variation after 42°C for Enterobacterales and 40°C for Pseudomonadales (Fig. S9).

To further assess potential limits on TPC evolution, we evaluated correlations among traits (Fig. 3A). We found that these correlations varied substantially at the Order level, likely indicating differences in patterns of TPC shape evolution across taxa. In Enterobacterales, increases in Topt were positively correlated with increases in performance peak (r = 0.28) and CTmax (r = 0.46), but less strongly with CTmin (r = 0.11). In contrast, increases in Topt in Pseudomonadales appeared to be accompanied by concomitant increases in CTmin (r = 0.52) and CTmax (r = 0.27), with virtually no change in performance peak (r = −0.01). We also saw a strong negative correlation between CTmin and CTmax in Pseudomonadales (r = −0.44).

The slope of performance decrease above Topt has been used as a proxy for the fitness costs of supraoptimal temperatures for microorganisms in marine environments. In those systems, a lower performance peak was predicted to be positively selected in highly variable environments to maximize thermal tolerance at the expense of maximum growth (Anderson et al. 2021). By definition, the slope above Topt (*b_T>Topt_*) will always be negative; the closer *b_T>Topt_* is to 0, the more robust growth will be with respect to increases in temperature. To maintain a given slope, if the maximum performance increases, the difference between CTmax and Topt must increase by the same factor. In our data set, this is not the case; as the performance peak increases, *b_T>Topt_* becomes steeper, with a consistent allometry (*b_T>Topt_* decreases 0.071 for each unit increase in AUCmax; Adj. R2= 0.72) across environments (Fig. S10).

To intuit how this slope is constrained, we obtained the maximum (gentlest or closest to 0) slope possible across values of AUCmax as the maximum distance between CTmax and Topt (Maximum CTmax observed - Minimum Topt observed) divided by AUCmax (max(*m_b/AUCmax_*)= −0.028). This slope becomes steeper as AUCmax increases but it is a gentler decline than our observed data (Fig. 3B). We then compared our results to slopes obtained from randomizing the thermal traits (mean(*m_b/AUCmax_*) = −0.101), and while our values fall within random expectations, they are significantly closer to the trend of “Gentlest slopes” than it would have been expected by chance (Fig. 3B, bottom panel; t = −14.368, df = 264, p-value < 2.2e-16). That is, observed slopes across taxa are close to the least steep value expected given the distribution of CTmax and Topt in our data (Fig. 3A).

The trend of increasing slopes with performance peaks could also be an artifact of fitting increasingly concave TPCs. This would lead to smaller ranges between CTmax and Topt values with performance peak, and a reduced variability. After binning the AUCmax values into quartiles for Pseudomonadales and Enterobacterales, we see that although variability in the CTmax-Topt range decreases with performance peaks, its width is relatively constant for Enterobacterales, and it increases for Pseudomonadales (Fig. S11).

The CTmax-Topt range also increases across CTmax quartiles for both taxa (Fig. S11), meaning that, as CTmax increases, Topt does not scale at the same rate, in spite of having a similar overall variance. This could have significant fitness costs in thermally variable environments. Due to the curves’ concavity at higher temperatures, performance at the mean temperatures is greater than the average performance across temperatures (*f*(*T*^-^) > ^-^*f*(*T*)). This is predicted to favor an increase in the difference between Topt and the mean annual temperature (MAT) in thermally variable climates to minimize the likelihood of fitness drops at extremely high temperatures, at the expense of a lower performance at suboptimal temperatures (Amarasekare and Johnson 2017; Martin and Huey 2008). We tested this by comparing (Topt - MAT) within colder and warmer climate profiles and observed contrasting trends: the difference increases with variability in cold environments, in agreement with the hypothesis based on Jensen’s inequality, but it decreases in hot environments (Fig. 3C).

### TPC peaks are biased towards high temperatures across sites

Limits in CTmax and Topt could result from common selective pressures rather than evolutionary constraints. That is, large values of CTmax might be uncommon because those temperatures are rare in the environment (low selective pressure). We analyzed the overlap between bacterial TPCs and yearly local temperature probability distributions and compared them to our TPCs (Fig. S12). Across environments, our TPCs matched the higher ends of the temperature distribution and missed growth at lower temperatures. Bacterial growth and activity throughout the entire year is unlikely. Instead, we predicted periods of higher activity based on estimates of plant growth season by site, but the overall patterns held (Fig. 4). We then calculated a metric of thermal fitness as a weighted average of AUC in growth across temperatures (*integrated thermal fitness;* ITC). Enterobacterales consistently showed higher ITC than Pseudomonadales across climate profiles, although the difference was smaller at hot and variable sites (Fig. 4, Inset).

**Figure 4:**
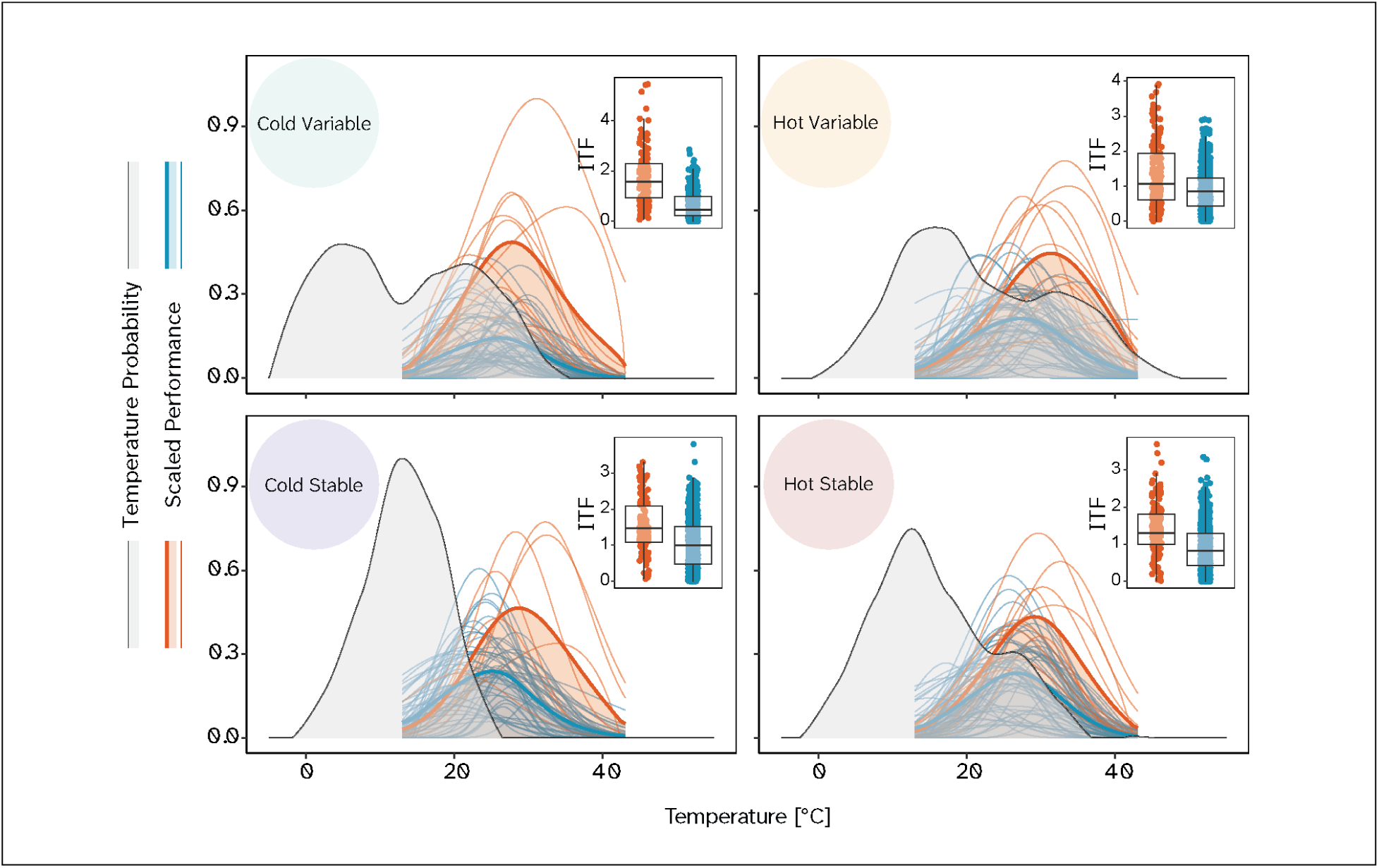
Overlaying the temperature probability by climate profile and the scaled performance by taxon shows differences in thermal fitness. We obtained a probability distribution of temperatures (light grey) by Climate Profile during the plant growth season and overlaid a scaled performance of each isolate and Order by temperature. Fine lines represent individual curves, and thick lines display average values for Enterobacterales (Orange) and Pseudomonadales (Blue). We then calculated the “Integrated Thermal Fitness” (ITF; inset) as the intersection area of each isolate curve with the temperature probability distribution of its climate profile.

Even when correcting for growth season, bacteria in hot climate profiles, especially those more variable, experience supraoptimal temperatures for a significant period (T > Topt = 22.0% for Enterobacterales and 30.6% for Pseudomonadales). At these sites, temperatures exceed the calculated CTmax by 0.44% and 2.74% of the time for Enterobacterales and Pseudomonadales, respectively (Fig. 4).

## Discussion

While it is expected that bacteria readily adapt to local conditions due to rapid growth rates and large population sizes, our results suggest that the evolution of thermal traits of copiotrophic Gammaproteobacteria is shaped more by phylogenetic and physiological constraints than present-day bioclimatic variables. In particular, Topt and CTmax showed little variation in our collection, and we observed a clear decline in overall growth at temperatures above 40°C. These upper growth limits are consistent with a mesophile–thermophile gap, with overall bacterial growth rates declining around 50°C (Corkrey et al. 2016) before increasing again due to thermophile taxa. Our data suggests that these constraints on upper thermal limits might already be costly, given that temperatures in hot-variable environments already exceed our thermal maxima at an average of one week a year, and in hot-variable environments they exceed the thermal optima between 30-40% of the time.

Phylogenetic signal in performance peak aligned with previously described and conserved metabolic strategies (Vila et al. 2023; Gralka et al. 2023), while shallower conservatism of thermal optima likely reflected genus-specific life-history traits. Although both thermal limits (CTmin and CTmax) lacked phylogenetic signal, we argue that this reflects different processes. The limited variation in growth at high temperatures (Fig. 3, Fig. S9) and a weak correlation with mean environmental temperatures suggest strong constraints on CTmax evolution, leading to conservation deeper than the maximum branch length in our tree (Fig. 5). In comparison, CTmin Fwas highly labile, with 93.8% of its residual variance unexplained by phylogeny. These findings are partially in agreement with previous evidence of higher evolutionary rates of CTmin compared to CTmax (Ørsted et al. 2022; Deutsch et al. 2008), but the little response to environmental conditions, and the fact that CTmin is much larger than the minimum temperatures encountered in all environments, indicate that the evolution of CTmin might not be strongly selected upon in these environments: for instance, these species might be dormant at low temperatures as it is common for other mesophiles (Lennon and Jones 2011; Du et al. 2007). However, these conclusions should be taken with caution: unlike other traits, estimation of CTmin is highly noisy due to the difficulties of measuring bacterial growth at very low temperatures.

**Figure 5:**
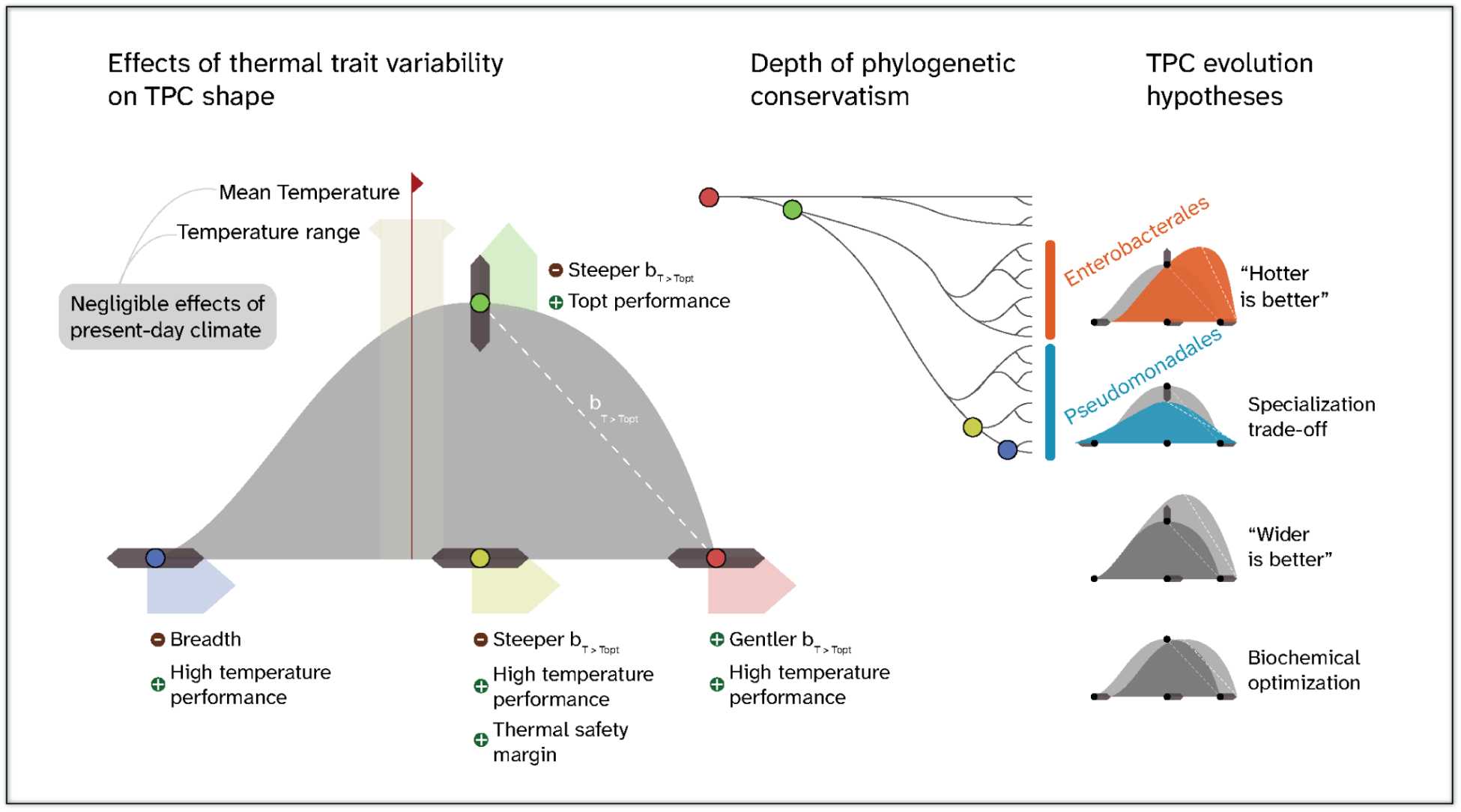
The differential conservatism of thermal traits reveals potential trade-offs in the TPC evolution of soil copiotrophs. This summary figure outlines the effects of increasing the value of different thermal traits on TPC shape, and their potential impacts on fitness in warming environments. Depth differences of phylogenetic conservatism suggest that TPC evolution is constrained in the direction and magnitude of change possible. Within this space of possible TPC shapes, Enterobacterales and Pseudomonadales exhibit different trait correlations, which could be explained by alternative models of TPC evolution.

In terms of evolution at higher temperatures, we expected that warmer environments would favor increases in Topt and CTmax, and that greater thermal variability would increase the overall range between CTmin and CTmax, as well as Topt, to extend the thermal safety margin. Instead, soil bioclimatic variables known to shape the TPCs of terrestrial ectotherms (Amarasekare and Johnson 2017; Ørsted et al. 2022; Deutsch et al. 2008) showed weak, clade-specific correlations with thermal limits and optima (Fig. S13). CTmax and Topt were significantly associated with thermal variability, but this association explained a minimal amount of variation. Other factors, such as ecological interactions or site-specific soil physicochemical properties, could play a greater role in shaping the TPC of these bacteria. For example, pH has been observed to dominate over temperature in the composition and diversity of soil microbial communities (Lauber et al. 2009), but the interaction between these environmental factors is not well understood. We need to evaluate physiological interactions across different environmental conditions and measure bacterial physiology in environments with enough complexity to understand the factors shaping bacterial thermal traits.

Our data allows us to evaluate potential constraints and relationships between different thermal traits. For instance, we can evaluate competing hypotheses for the TPC evolution of ectotherms in warming environments (Malusare et al. 2023). These hypotheses differ on the relative influence of thermodynamic, physiological, and ecological factors on the variation of height and breadth of the TPC. The “Hotter is Better” hypothesis posits that the maximum performance of curves is thermodynamically constrained, and thus that performance peaks and thermal optima are positively correlated, whereas alternative hypotheses propose that physiological adaptations can lead to equivalent performance peaks along the thermal spectrum (Angilletta et al. 2010).

Other models introduced specialization trade-offs to explain simultaneous shifts in the height and width of TPCs (Izem and Kingsolver 2005), and have been successfully associated with estimations of ecological niche in phytoplankton (Krinos et al. 2025). Within our collection, TPCs of Enterobacterales aligned with the “Hotter is Better” hypothesis, with a positive Topt-peak performance correlation, where performance peak is favored over risk minimization, while the lower and less skewed TPCs of Pseudomonadales appear to be favoring a temperature-generalist strategy, with lower temperature optima and a negative correlation between CTmin and CTmax (Fig. 5).

Global temperature rises are accompanied by increased thermal variability. Adaptation to these scenarios is expected to involve increases in Topt, CTmax, and thermal safety margin to minimize catastrophic fitness losses during extended heat anomalies (Amarasekare and Johnson 2017; Martin and Huey 2008). Because TPCs are concave at high temperatures, it is expected that the Topt should exceed the mean annual temperature to maximize performance averaged over time (Amarasekare and Johnson 2017). Instead, we found that while hot variable environments have higher average CTmax, they also have smaller safety margins, indicating potential maladaptation due to constraints in the evolution of thermal traits.

Despite constraints, we observed possible evidence of evolutionary optimization when comparing the steepness of supraoptimal temperature slopes. All else being equal, minimizing the steepness of these slopes provides an advantage, especially in hot variable environments where temperatures are likely to exceed the thermal optima frequently (Amarasekare and Johnson 2017; Martin and Huey 2008). We observed a remarkably conserved relationship between this steepness and the maximum performance (Fig. 3B). By randomizing the thermal traits, we showed that these slopes are generally less steep than those expected by chance. But they also reflect how constraints in some traits can affect the overall geometry of the TPC: given the robustness of CTmax and Topt, increasing peak performance necessarily entails an increase in the steepness of this supraoptimal slope. More detailed curves, fitted with the same model, are necessary to carefully compare and evaluate the evolution and constraints on the curvature of these thermal responses.

It is unclear how much we can extrapolate from laboratory physiological assays to the field. The selective pressure of extreme heat events not only depends on their recurrence and intensity, but also on correlated stressors, such as drought. Consequently, the upper thermal limits in natural settings are likely lower than those estimated in liquid culture. Bacterial populations interact within a dynamic biophysical environment, and the TPC shape of a given species can depend on interspecific interactions (Westley et al. 2024) and nutritional context (Hutchins and Tagliabue 2024). Other ecological factors, such as the production of biofilms that buffer environmental variability (Yin et al. 2019), the horizontal acquisition of stress-resistance genes (Shi and Thakur 2023; Rzymski et al. 2024), or the dispersal to endothermic hosts (Khakisahneh et al. 2020), could alleviate the pressure for growth at high and variable temperatures. Finally, extreme temperatures might favor life-history strategies different from those represented by our copiotrophic taxa. This is the case for stress-resistant life-history strategies, which are often characterized by low growth rates and high resource allocation towards cell maintenance (Malik et al. 2020; Piton et al. 2023; Pold et al. 2015).

Our work here focused on fast-growing bacteria using a labile, energy-rich resource to expose constraints on thermal performance at high metabolic rates. Overall, we highlight potential limits to TPC evolution and the importance of analyzing different thermal traits and their interactions. Using a comparative physiology approach, we provide support for the idea that each of these traits is shaped by different evolutionary processes and physiological constraints, leading to the evolution of potential trade-offs among thermal strategies. Our data provides support for constrained evolution of thermal maxima in these taxa and suggests that increasing temperatures and variability could limit copiotrophic performance.

## Acknowledgements

MRG was partially supported by the John Templeton Foundation grant 63455. AF was partially funded by the UCI 2022 Miguel Vélez Scholarship. We obtained soil samples at fifteen sites across fourteen reserves of the University of California Natural Reserve System: Bodega Marine Reserve (10.21973/N32Q05), Boyd Deep Canyon Desert Research Center (10.21973/N3V66D), Elliott Chaparral Reserve (10.21973/N36H23), James San Jacinto Mountains Reserve (10.21973/N3KQ0T), Jepson Prairie Reserve (10.21973/N3D082), McLaughlin Natural Reserve (10.21973/N3W08D), Merced Vernal Pools and Grassland Reserve (10.21973/N3FT0M), Point Reyes Field Station, Quail Ridge Reserve (10.21973/N30T0K), Yosemite Field Station (10.21973/N3V36C), Steele/Burnand Anza-Borrego Desert Research Center (10.21973/N3Q94F), Sierra Nevada Aquatic Research Laboratory (10.21973/N3966F), Valentine Camp (10.21973/N3JQ0H), White Mountain Research Center (10.21973/N3KM2J).

## Statement of authorship

AIF and MR-G designed the study with input from CCS. AIF and AH-T collected data. AIF and MR-G analyzed data with help of AH-T. AIF wrote the first draft of the manuscript with substantial input from MR-G. All authors contributed to subsequent drafts.

## Data accessibility statement

All growth data and code can be found in our GitHub repository https://github.com/mrebolleda/Paper_PhylogeneticThermalConstraints (a static copy will be made available upon publication). Raw sequencing reads are available in the NCBI Sequence Read Archive (SRA) under BioProject accession **PRJNA1414602**. We partially processed the genomic data using a KBase Narrative (https://doi.org/10.25982/200311.171/3014786).

