## Supplementary material for "Conserved upper thermal limits and small safety margins in soil copiotrophic bacteria": 260206_Supplementary_figures_and_methods

**Supplementary Figures and Methods for "Conserved upper thermal limits and small safety margins in soil copiotrophic bacteria"**

| Supplemental Figure | Script | Name in Script |
|---|---|---|
| 1 | 251001_precipitation_db_comparison.R | S1 |
| 2 | 251029_TCS_TPCs.R | S2 |
| 3 | 251029_TCS_TPCs.R | S3 |
| 4 | 251029_TCS_TPCs.R | S4 |
| 5 | 251029_TCS_TPCs.R | S5 |
| 6 | 251029_TCS_figures | S6 |
| 7 | 251029_Environmental_data_processing | S7 |
| 8 | 251029_TCS_figures | S8.left and S8.right |
| 9 | 251029_TCS_figures | S9.top and S9.bot |
| 10 | 251029_TCS_figures | S10 |
| 11 | 251029_TCS_figures | S11 |
| 12 | 251029_TCS_figures | S12 |
| 13 | 251029_TCS_figures | S13.top.left, S13.top.center, S13.top.right, S13.bot.left, S13.bot.center, S13.bot.right |
| 14 | 251029_TCS_figures | S14B.left and S14B.right |
| 15 | 251029_TCS_figures | S15.bot |
| 16 | 251029_TCS_figures | S16A, S16B, and S16C |
| 17 | 251029_TCS_figures | S17 |

### *Whole Genome Sequencing and Annotation*

We extracted the genomic DNA of 400 randomly selected isolates using a Quick-DNA Microprep Kit (Zymo Research D3020) according to the manufacturer's protocol. We then submitted the extracted gDNA samples for short-read Illumina sequencing (200 Mbp) at SeqCoast Genomics (Portsmouth, NH, USA). After preprocessing the sequences using Trimmommatic (Bolger et al. 2014), we assembled the genomes using SPADES (Bankevich et al. 2012) and checked the quality of each assembly using QUAST (Gurevich et al. 2013). We processed the genome assemblies using a KBase (v1.4.0) pipeline (Allen et al. 2017; Arkin et al. 2018). Briefly, we used DRAM (v0.1.2) with default settings to annotate the genome assemblies. We then evaluated genome quality and possible contamination levels using CheckM (v1.0.18) (Parks et al. 2015) and retained genomes with completeness above 98% and contamination below 5% (n = 354), following the authors' guidelines. We then obtained taxonomic assignments for all remaining isolates using the Genome Taxonomy Database tool GTDB-Tk (v2.3.2, database version r214) (Chaumeil et al. 2019). We constructed a phylogenetic tree using the tool SpeciesTree (v2.2.0), which identifies a set of 49 core universal genes from families within COG (Clusters of Orthologous Groups) in the uploaded sequences, aligns them with others from closely-related public KBase genomes, concatenates them, and finally reconstructs an Approximately-Maximum-Likelihood phylogenetic tree using FastTree (v2.2) (Price et al. 2010). We then trimmed the tree (using Trim SpeciesTree to GenomeSet- v1.4.0), retaining only tips within our collection with measured thermal performance.

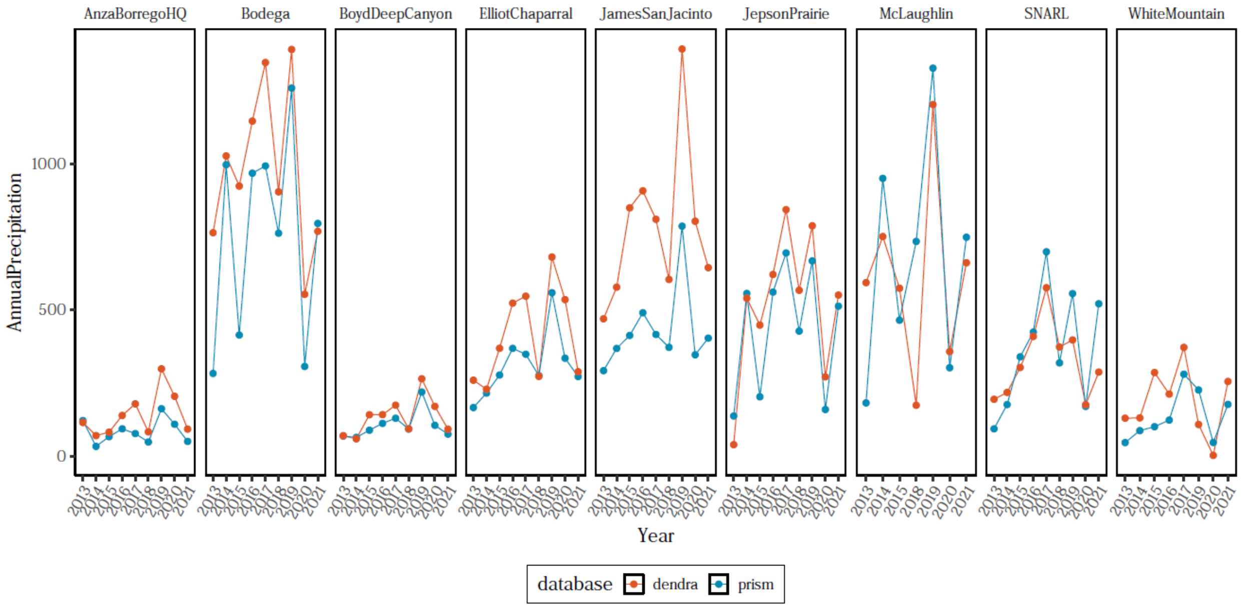

**Supplementary Figure 1: Comparison of annual precipitation measurements between databases.** Sites with more than five years of data available in both sources generally showed consistent values across databases (except for James San Jacinto), giving us confidence in their comparability. As a result, we used the more complete PRISM time series as our precipitation database for all sites.

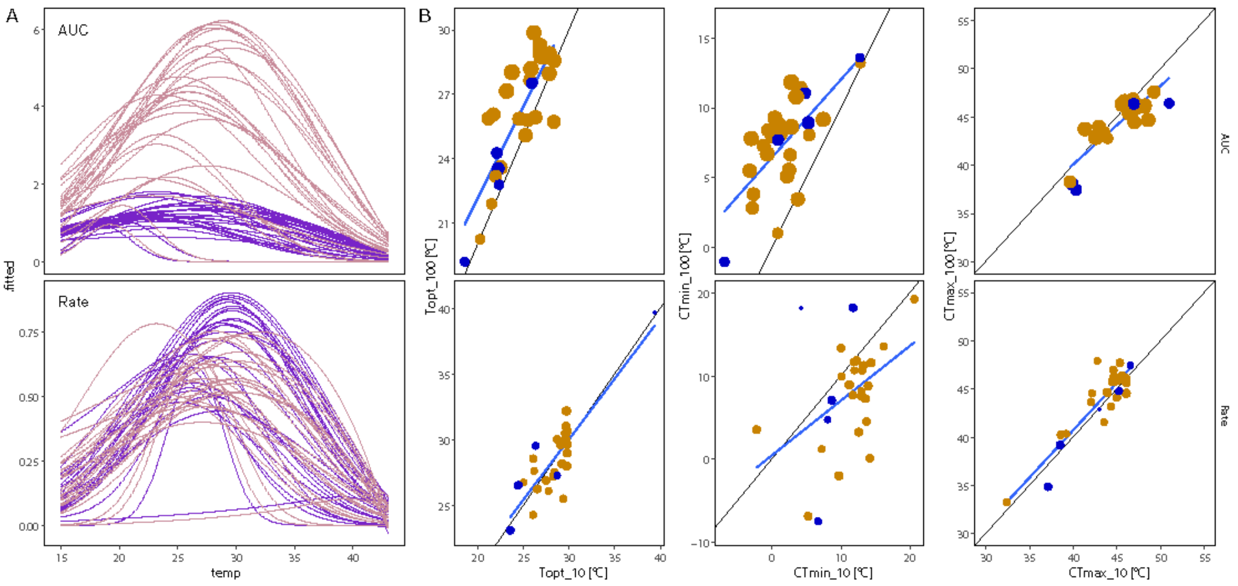

**Supplementary Figure 2: Changes in carbon concentration do not impact key TPC thermal traits significantly across metrics.** (A) We fitted TPCs of a subset of 28 isolates after growth at 11 different temperatures in M9-glucose media with 0.007 (purple) and 0.07 (pink) M-C final carbon concentration. (B) We then compared the TPC traits between the high (0.07 M-C = 100) and low (0.007 M-C = 10) carbon concentrations. The black line indicates a theoretical 1:1 relationship. The points are colored by Order (Yellow = Enterobacterales, Blue = Pseudomonadales) and sized by peak performance.

### *Growth curve fitting*

We used the R package *gcplyr (Blazanin 2023)* to fit models to our growth curves and to obtain the main growth parameters for each measurement. We took several steps to ensure data reliability and minimizing effects of fitting artifacts. Using the function *movavg*, we smoothed the raw absorbance data for all curves by calculating a 21-time-point moving average window (3.5 hs) to minimize the effects of noisy measurements, particularly prevalent at extreme temperatures. Subsequently, we calculated the derivative of absorbance using the function *calc_deriv* over 29 smoothed absorbance values (4.83 hs), a time interval that approaches the median length of the exponential phase in our cultures across temperatures (~5.61 hs; Fig. S4). To calculate the growth parameters, we excluded absorbances below 0.01 and time values under 3 hours from the analysis, as the presence of outliers strongly affected the derivatives of low-absorbance values. Finally, we manually curated the database by examining the fits of each isolate and temperature, removing evident outliers (curves with mismatched maximum growth rates and odmax values, Fig. S5) and previously documented contaminations.

### *Thermal performance curve (TPC) fitting*

We applied a filtering step before the model AICc weighting to remove those models that fitted biologically unrealistic curves, mainly characterized by a very narrow breadth and exaggerated maximum value, by applying a cutoff value of breadth > 5°C and the maximum parameter estimates obtained by growth curve fitting, after visual inspection of individual curves (Fig. S7). When computing the weighted averages, we omitted instances of infinite values. After plotting the average models, we removed those isolates with unrealistic results and kept a total of 244 for further analysis.

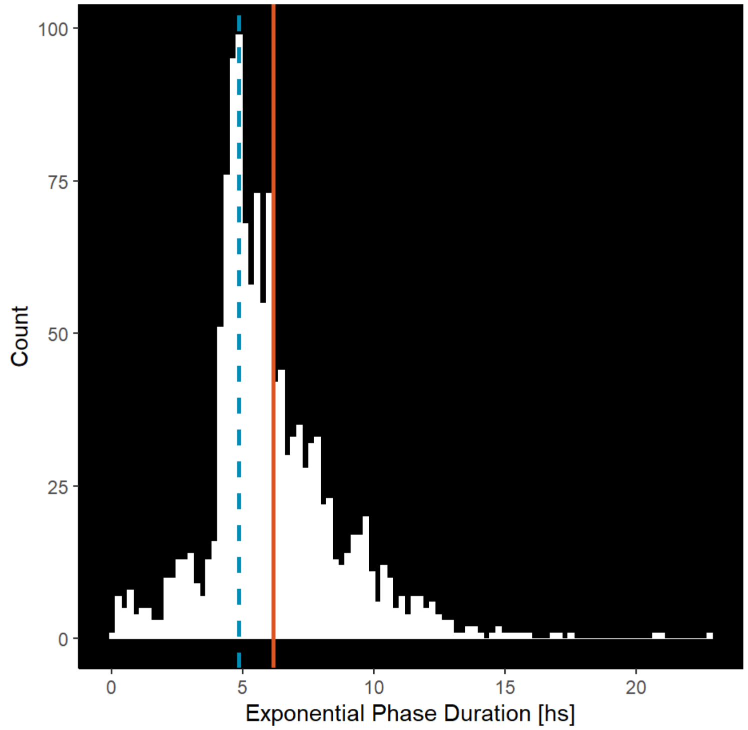

**Supplementary Figure 3: Selection of a sliding window for calculating the maximum growth rate.** We plotted the distribution of Exponential Phase Duration (EPD) across temperatures for growth curves with valid estimates (0 hs < EPD > 24 hs). The window to calculate the maximum derivative should be narrow enough to avoid including the lag or the stationary growth phase, but wide enough to be robust to noise. Because of this, we selected a window (29 time points, 4.83 hs; dashed blue line) that falls roughly at 0.85x of the mean EPD (5.61 hs; solid orange line).

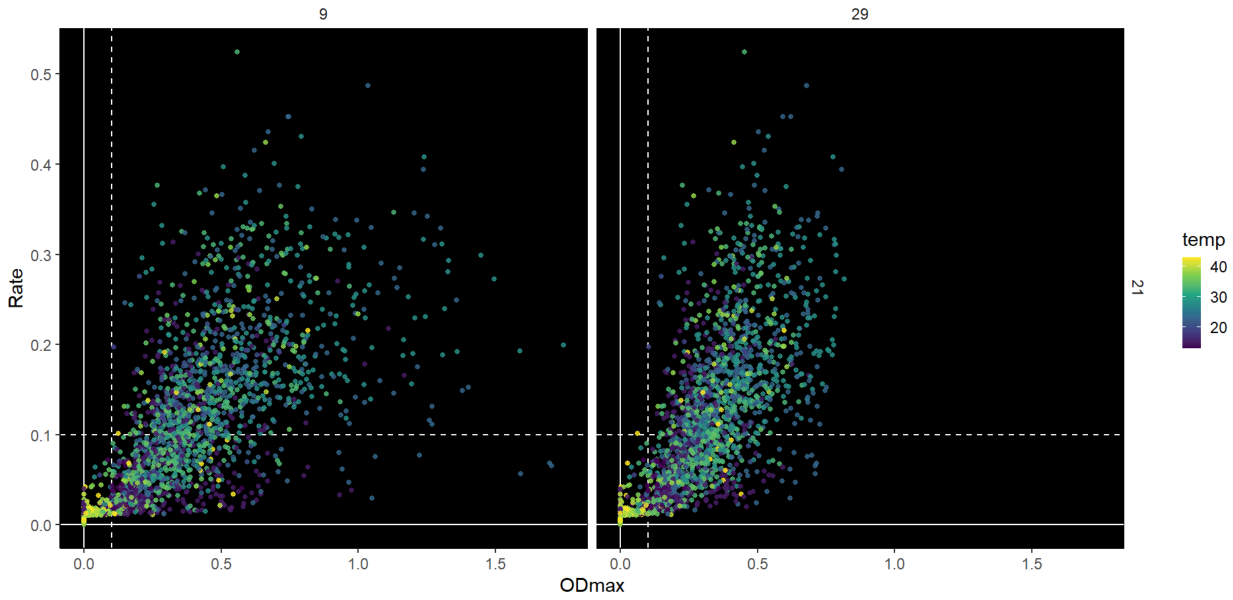

**Supplementary Figure 4: Comparison of growth curve parameters obtained using different sliding windows for absorbance and maximum growth rate.** We plotted the maximum growth rate (Rate) and maximum absorbance (ODmax) of our collection, faceted by the combinations of absorbance moving average (rows) and maximum growth rate window (columns), on a scale of time points used (29 = 29 time points = 4.83 hs). Points, colored by experimental temperature, represent average values for each isolate.

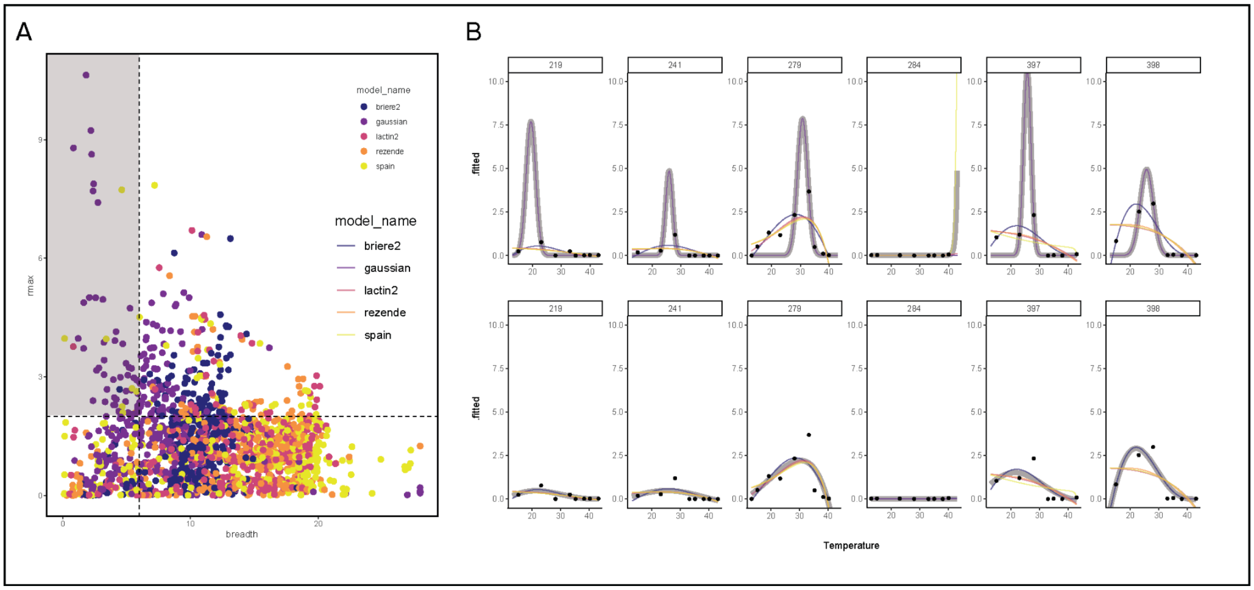

**Supplementary Figure 5: Removing well-fitting, unrealistic thermal performance curve models before calculating weighted averages.** (A) After fitting five different TPC models for each isolate, we generated a scatter plot of calculated peak performance (rmax) against breadth values, coloring the points by model. This allowed us to systematically identify potential artifacts without fully excluding otherwise well-fitting models. (B) We then examined specific cases (top) and removed those unrealistic models that would dominate the average fit (wider grey line) if included (bottom).

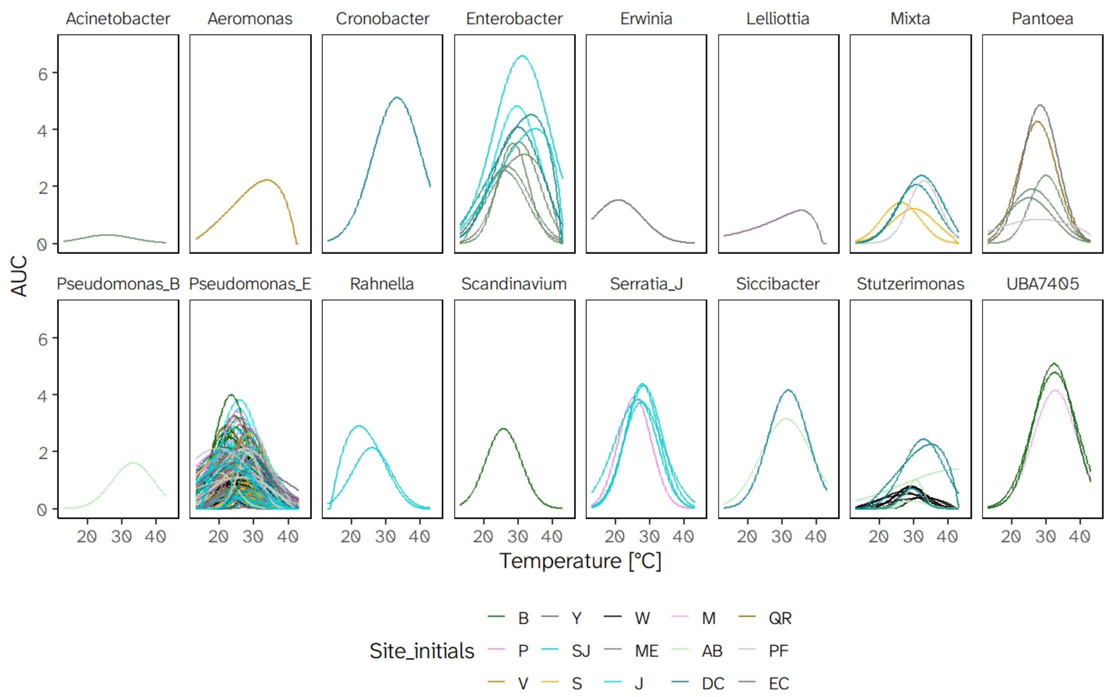

**Supplementary Figure 6: Thermal performance curves of AUC values grouped by Genus and colored by Site.** Site abbreviations are as follows: W - White Mountain Research Center, SJ - James San Jacinto Mountains Reserve, S - Sierra Nevada Aquatic Research Laboratory; V - Valentine Camp, Y - Yosemite Field Station, P - Point Reyes Field Station, B - Bodega Bay Research Center; AB - Steele/Burnand Anza-Borrego Desert Research Center Research Center, DC - Boyd Deep Canyon Desert Research Center, EC - Elliot Chaparral Reserve, QR - Quail Ridge Reserve, PF - Pinyon Flats; ME - Merced Vernal Pools and Grasslands Reserve, M - James McLaughlin Natural Reserve, J - Jepson Prairie Reserve.

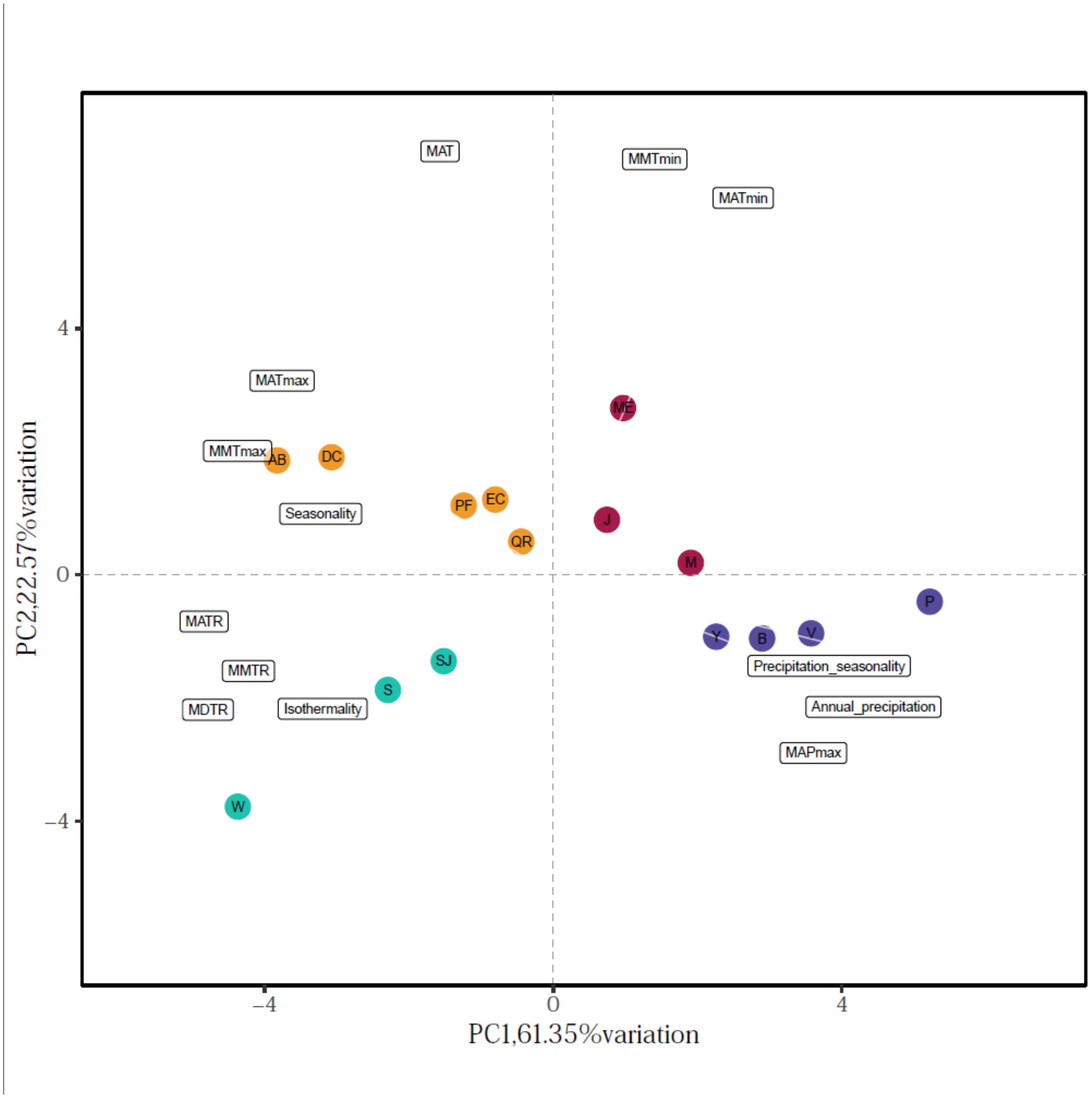

**Supplementary Figure 7: Principal Components of environmental variation, including precipitation variables.** (A) We obtained soil bioclimatic variables across sampling sites and performed Principal Component Analysis (PCA) to compress their variability into orthogonal components. The first two PCs explained a cumulative 83.92% of the variance across sites, compared to an 88% when excluding precipitation variables. Site abbreviations by temperature profile are (Cold Variable): W - White Mountain Research Center, SJ - James San Jacinto Mountains Reserve, S - Sierra Nevada Aquatic Research Laboratory; (Cold Stable): V - Valentine Camp, Y - Yosemite Field Station, P - Point Reyes Field Station, B - Bodega Bay Research Center; (Hot Variable): AB - Steele/Burnand Anza-Borrego Desert Research Center Research Center, DC - Boyd Deep Canyon Desert Research Center, EC - Elliot Chaparral Reserve, QR - Quail Ridge Reserve, PF - Pinyon Flats; (Hot Stable): ME - Merced Vernal Pools and Grasslands Reserve, M - James McLaughlin Natural Reserve, J - Jepson Prairie Reserve.

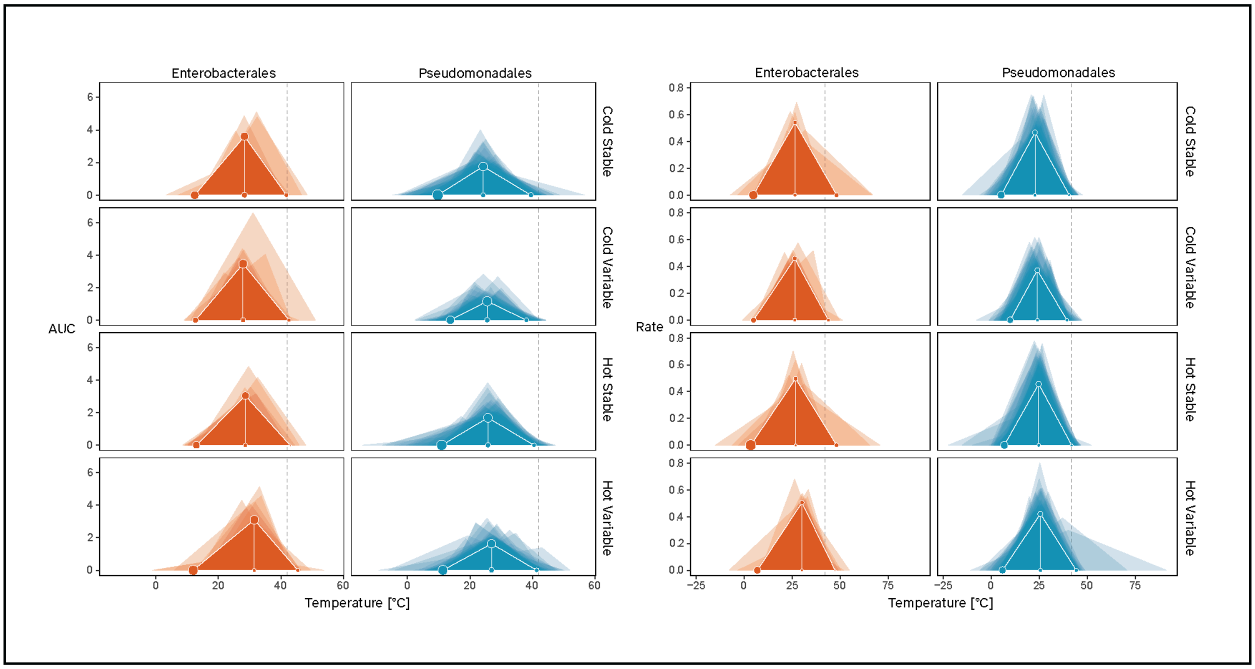

**Supplementary Figure 8: Simplified thermal performance curves for Enterobacterales (Orange) and Pseudomonadales (Blue) across climate profiles and growth metrics**. The main traits are represented by points (CTmin, CTmax, Topt, and Peak) and connected with straight lines to form a triangle for each curve. The dashed line indicates the upper thermal limit documented for mesophilic enzymes at 42°C. Translucent triangles represent individual isolates, and full triangles represent average values by taxon. Point size is proportional to the variability of the trait within that taxon, as in SD(trait)/Mean(trait).

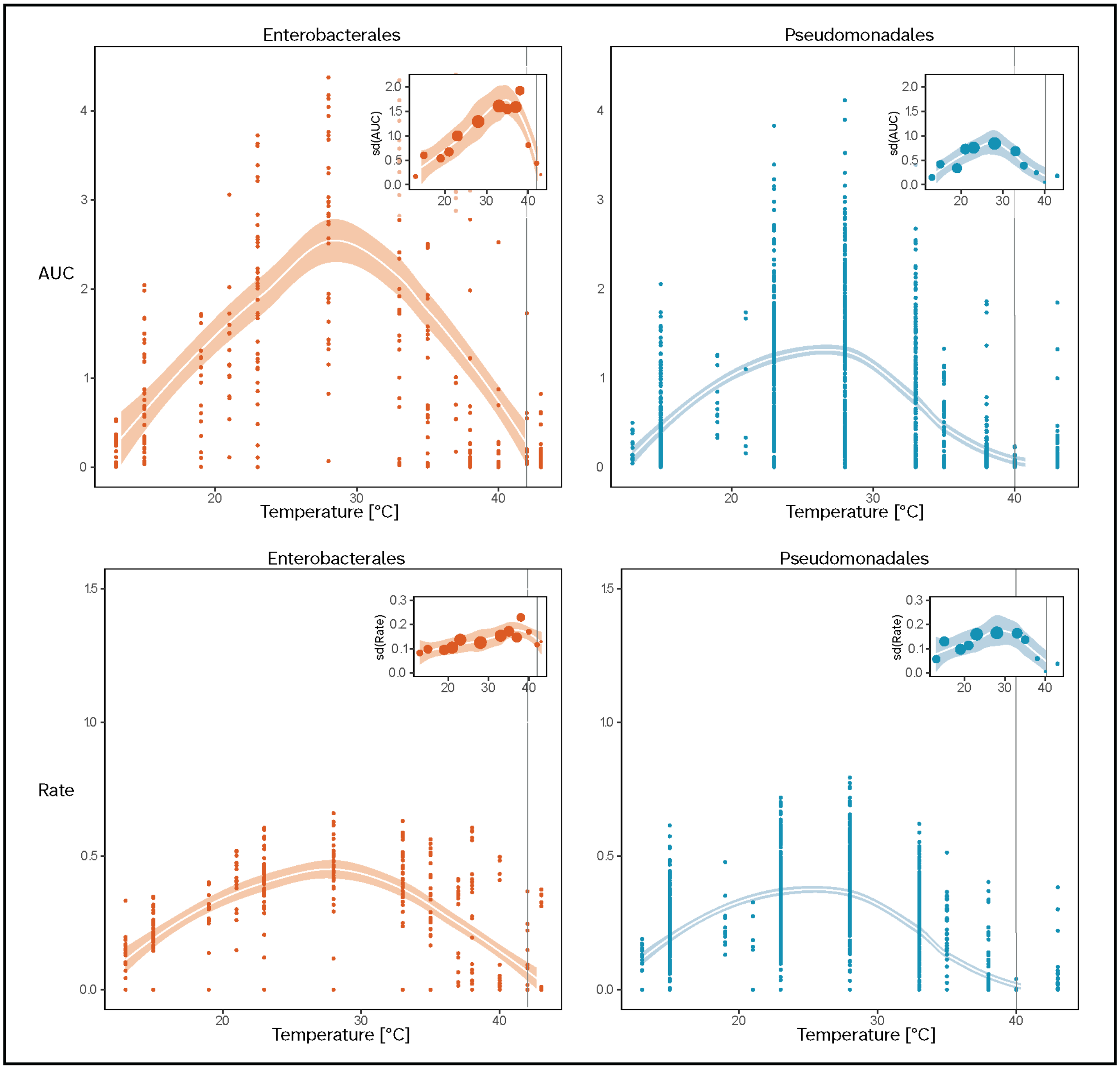

**Supplementary Figure 9: The distribution of growth curve parameters by temperature for the two major taxa in our collection**. The main panels show a scatterplot of the average growth values by isolate for Enterobacterales (Orange) and Pseudomonadales (Blue) by metric. The inset panels show the standard deviation of growth by measured temperature and Order. Shaded trendlines represent a 95% CI for each distribution. Grey vertical lines indicate the temperature at which both growth means and SD decrease below 25% of the peak.

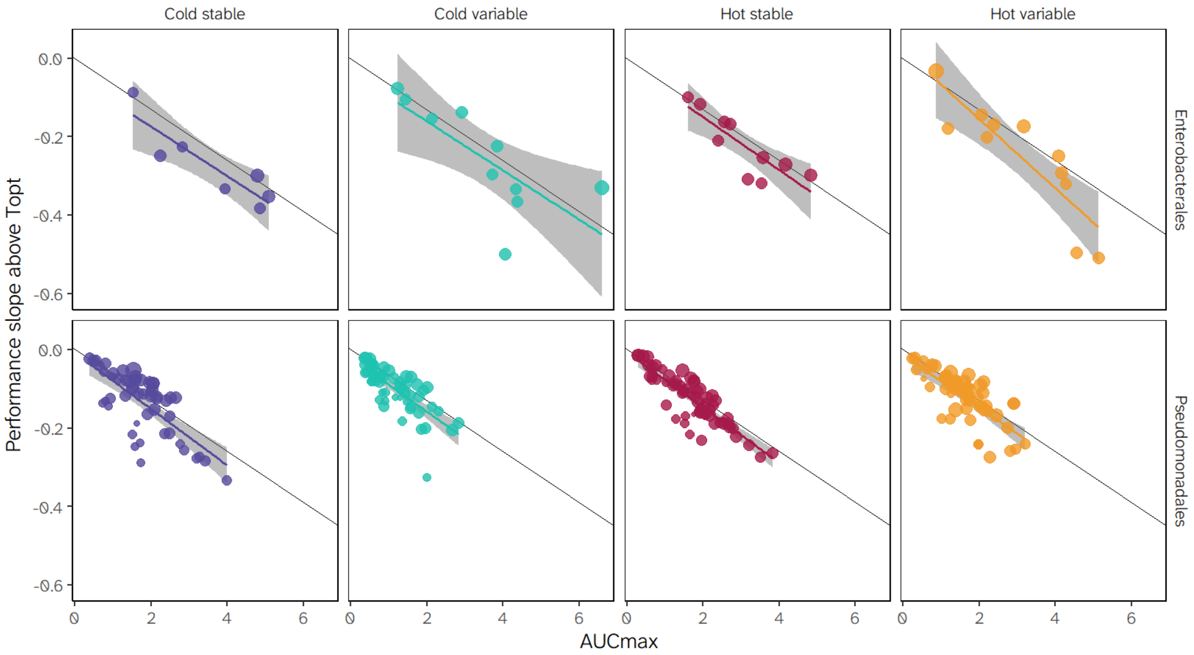

**Supplementary Figure 10: Changes in performance slopes above Topt are independent of climate profile and taxa.** We calculated the performance decrease slope above each simplified TPC's thermal optimum. Here, we show that the observed slopes become steeper with performance peaks, suggesting the apparent robustness of the upper thermal limit, and that this trend is generally maintained across taxa and climate profiles.

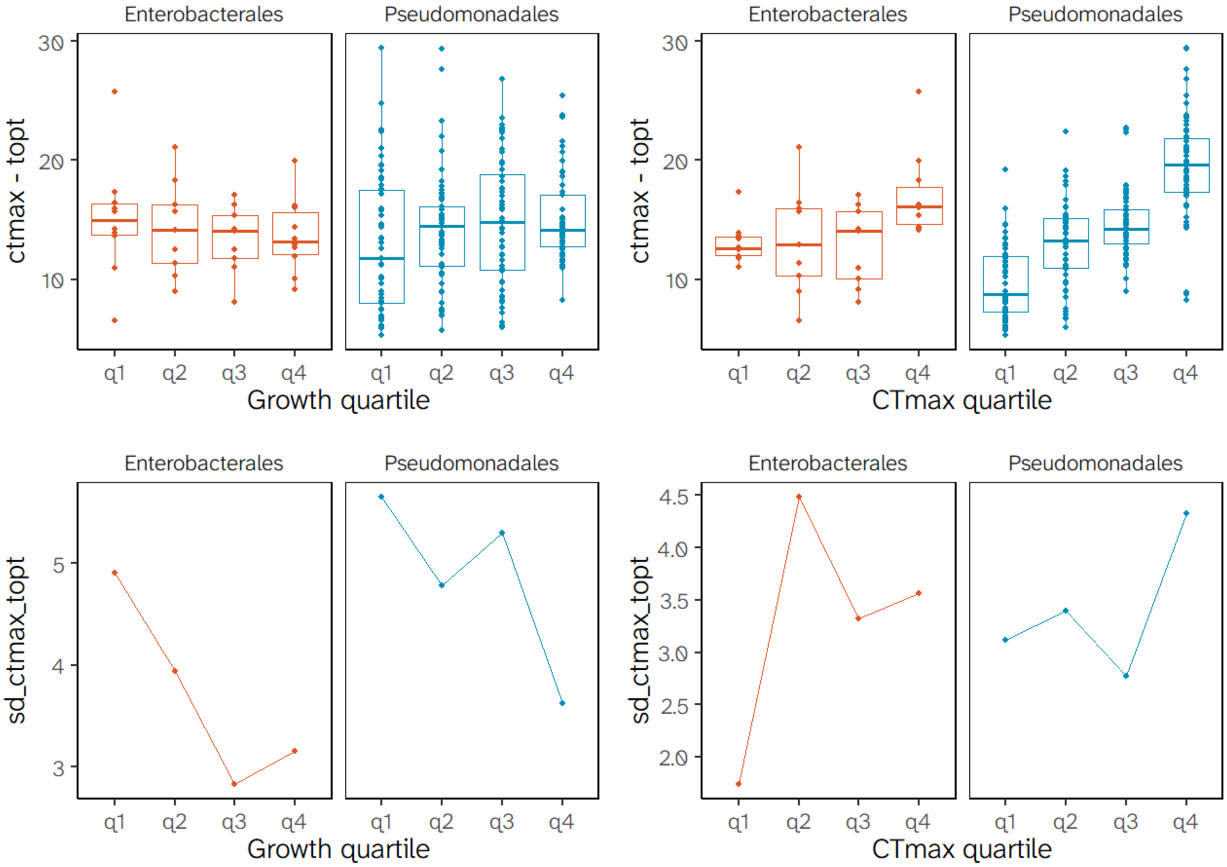

**Supplementary Figure 11: Changes in the mean and variability of the CTmax-Topt range.** We subset the isolates by Order into quartiles of either the performance peak (Growth; left) or CTmax (right). We show the changes in the range width (top) and standard deviation across quartiles.

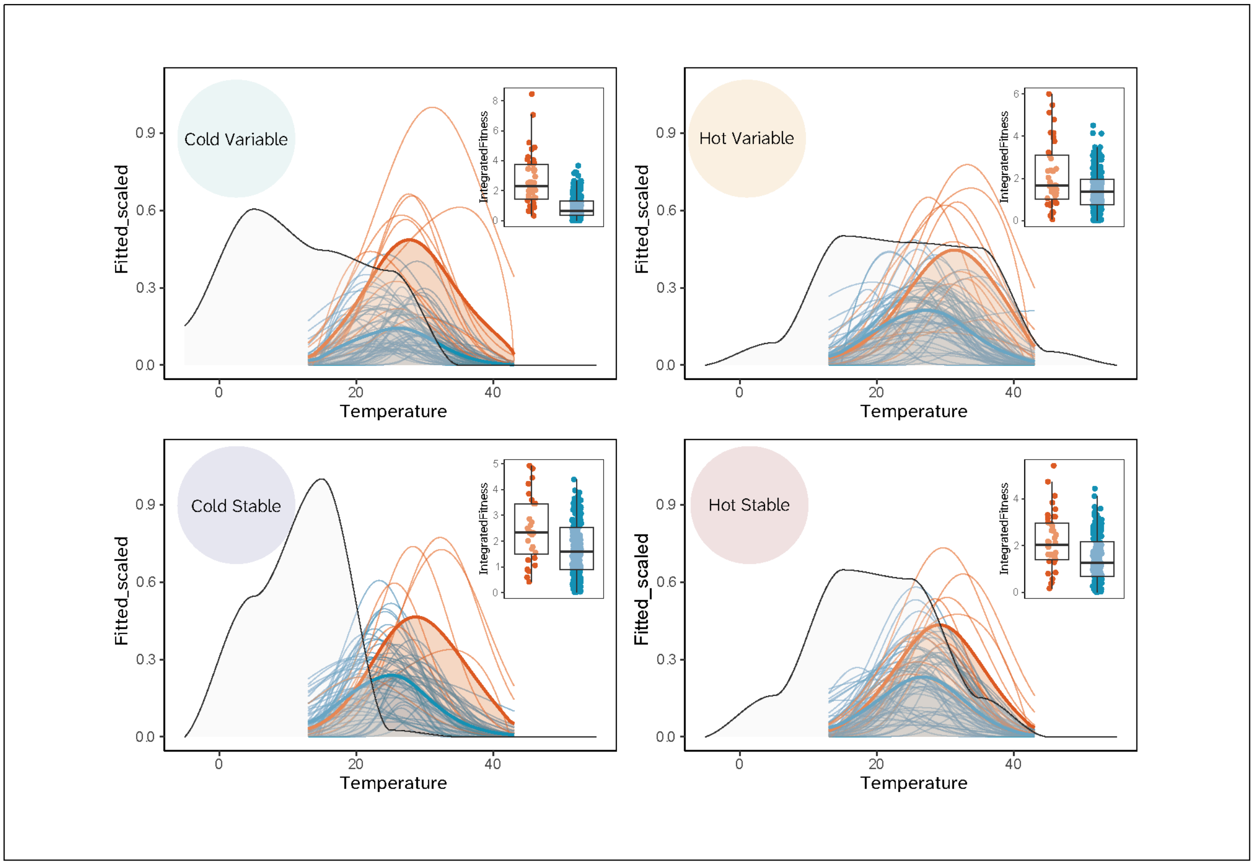

**Supplementary Figure 12: Overlapping the thermal performance curves and the yearly probability distribution of temperatures (light grey) by climate profile**. Fine lines represent individual curves, and thick lines display average values for Enterobacterales (Orange) and Pseudomonadales (Blue). Within each panel, the "Integrated Fitness" (inset) represents the area of intersection of individual curves with the temperature probability distribution.

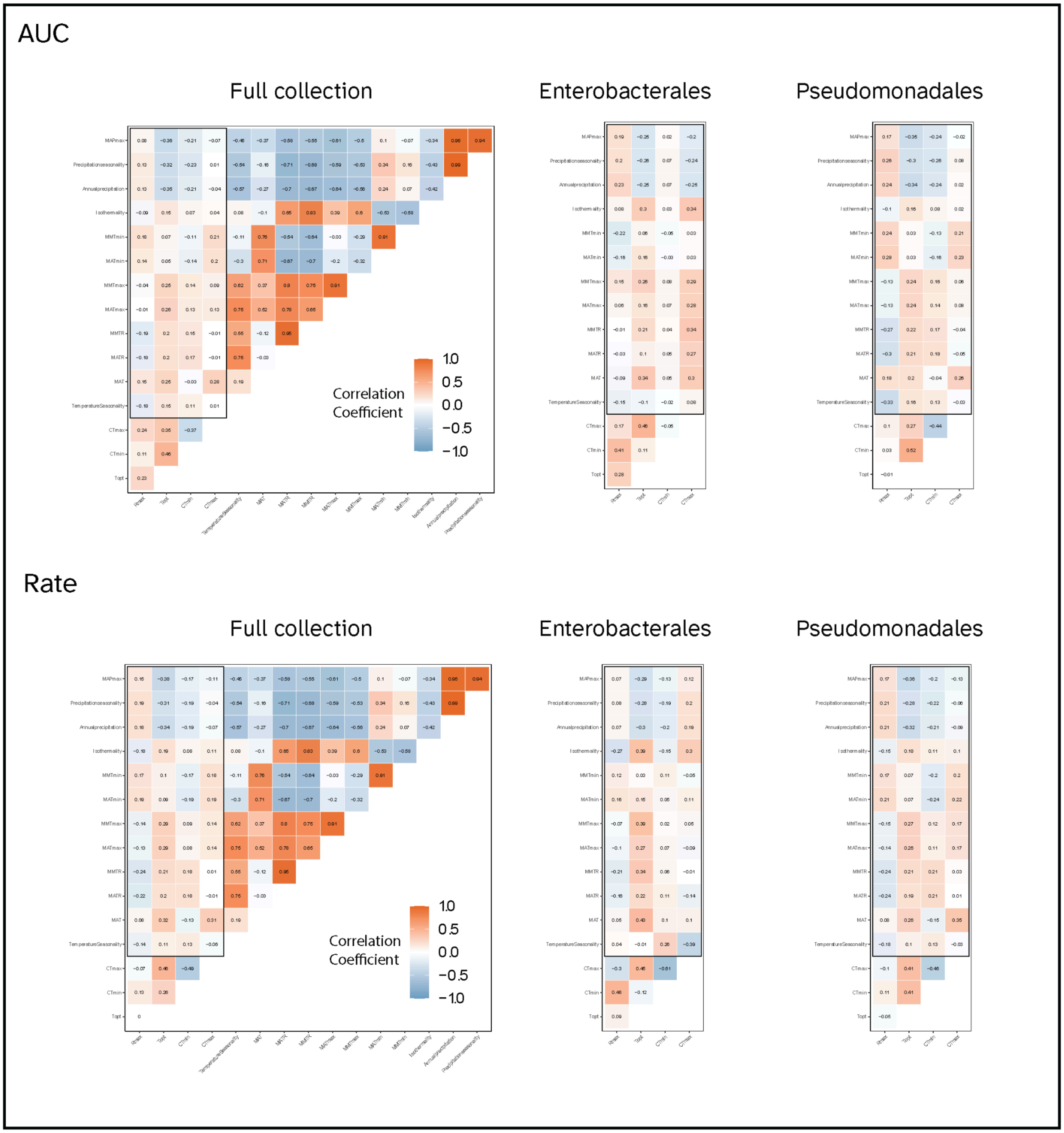

**Supplementary Figure 13: Linear correlation matrices between thermal traits and bioclimatic variables by growth metric**. From left to right, we first see the correlation matrix for the entire isolate collection with values colored as a gradient from negative (colder colors) to positive (warmer colors); the inner rectangle highlights the correlations between traits and bioclimatic variables, while the values outside of it are correlation coefficients between traits or between bioclimatic variables. We then see correlation matrices for Enterobacterales (center) and Pseudomonadales (right).

Rate

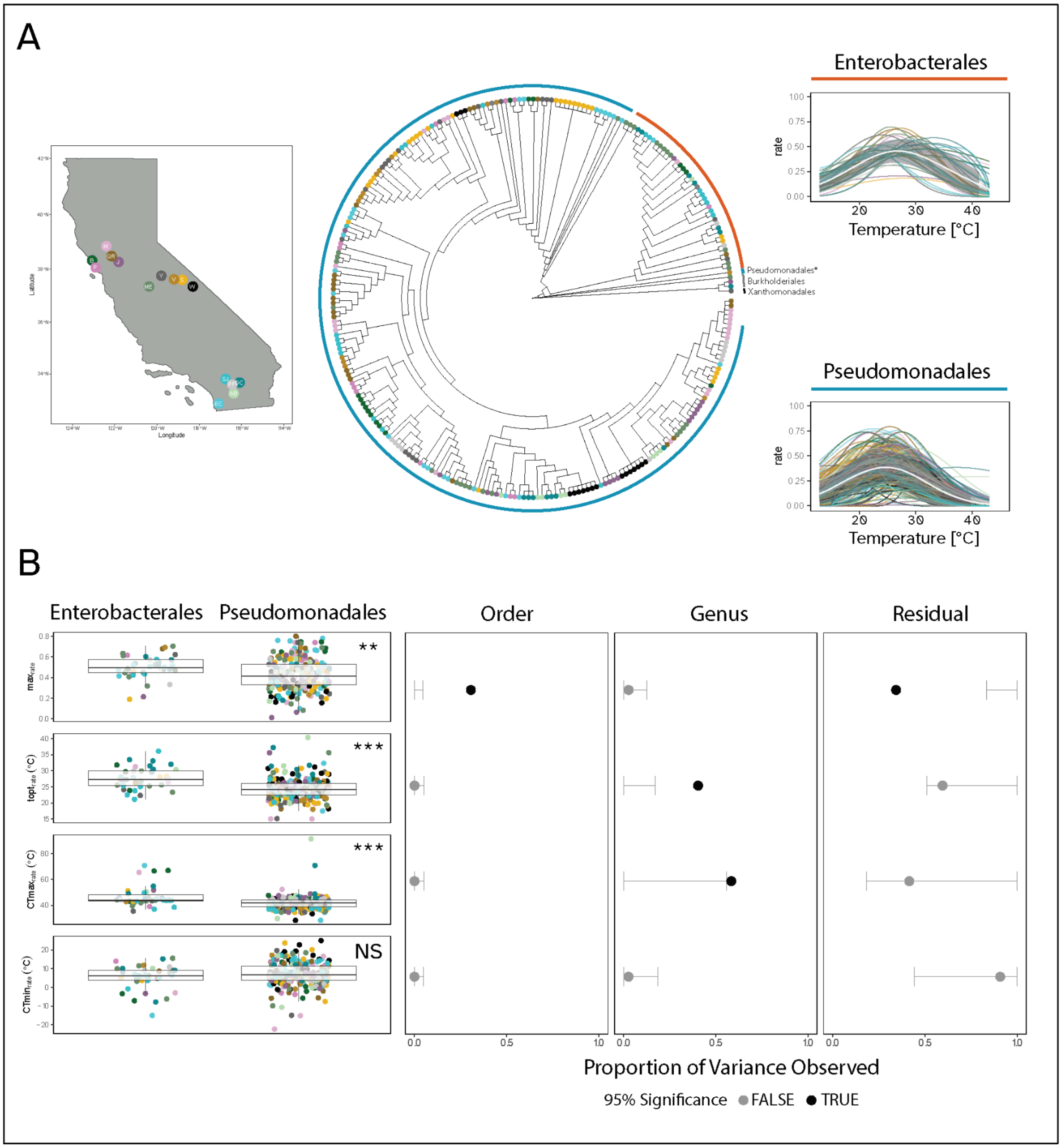

**Supplementary Figure 14: Thermal traits of growth rate thermal performance curves are differentially conserved in a collection of bacterial isolates from Californian topsoils. (A)** We collected topsoil samples from fifteen sites within the UCNRS and isolated bacteria at three temperatures (left). We sequenced a random sample of four hundred isolates and constructed a phylogeny (center). Points are colored to represent the sampling site, and the ring around the dendrogram is colored according to Order. The “*” signals a representative of the family Moraxellaceae, which consistently grouped outside of the main branch of Pseudomonadales. We then measured the thermal performance curves of the two major Orders (right). Shaded trendlines signal the mean SD of Rate across temperatures. **(B)** We compared the thermal traits between Enterobacterales and Pseudomonadales (left) using one-way

ANOVAs, with a 95% significance criterion (*** = $p < 0.001$, ** = $p < 0.01$, * = $p < 0.05$, NS = $p >= 0.05$), and we then compared the variance observed at each relevant taxonomic level to a confidence interval (CI) of variance expected after permuting trait values across the collection (nsim = 10000) (right). Black points outside the CI for the Order or Genus columns represent phylogenetic signal at that taxonomic level, while grey points represent the opposite. Site abbreviations are as follows: W - White Mountain Research Center, SJ - James San Jacinto Mountains Reserve, S - Sierra Nevada Aquatic Research Laboratory; V - Valentine Camp, Y - Yosemite Field Station, P - Point Reyes Field Station, B - Bodega Bay Research Center; AB - Steele/Burnand Anza-Borrego Desert Research Center Research Center, DC - Boyd Deep Canyon Desert Research Center, EC - Elliot Chaparral Reserve, QR - Quail Ridge Reserve, PF - Pinyon Flats; ME - Merced Vernal Pools and Grasslands Reserve, M - James McLaughlin Natural Reserve, J - Jepson Prairie Reserve.

Rate

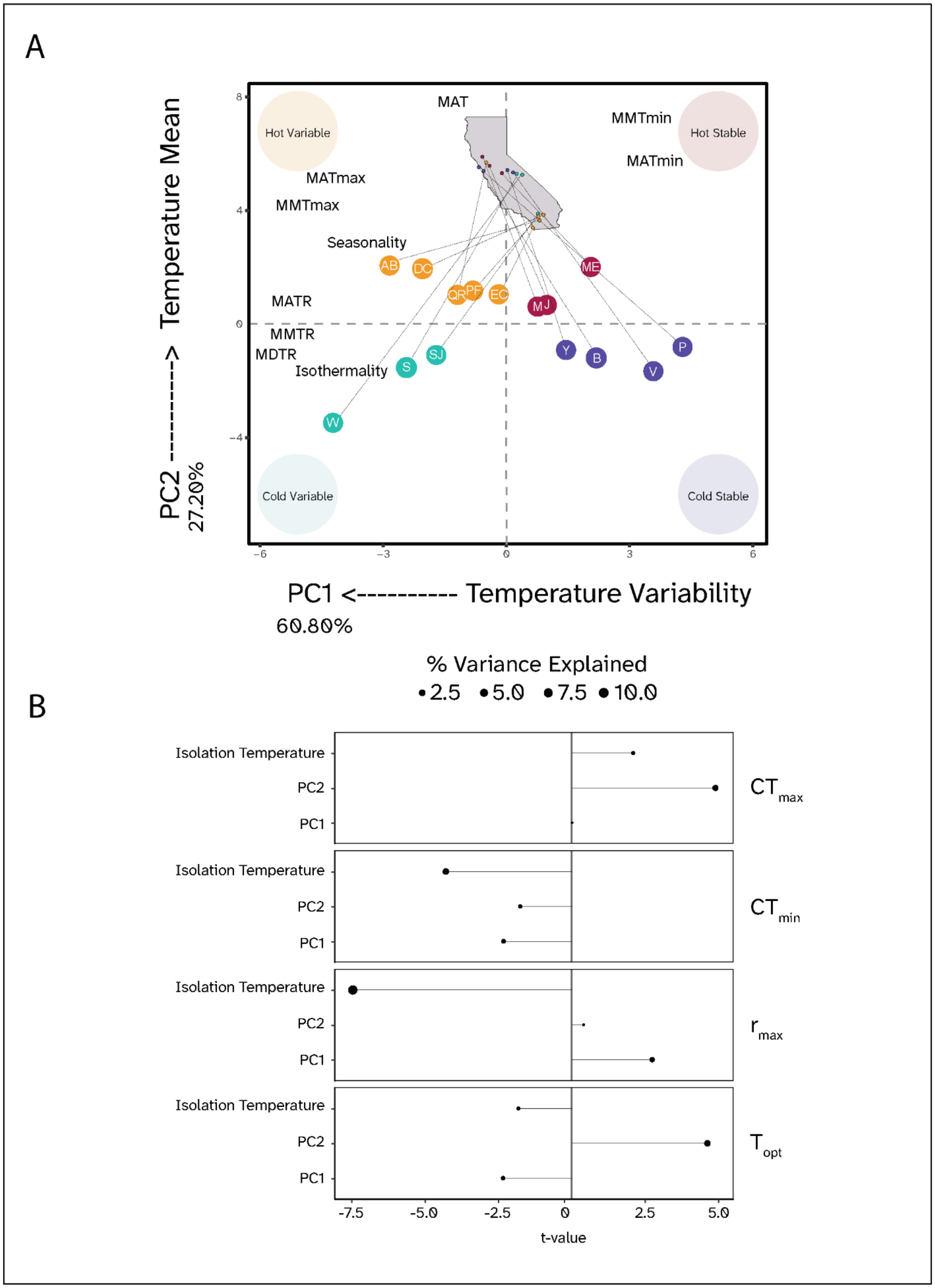

A
Hot Variable
Hot Stable
Cold Variable
Cold Stable
MAT
MMTmin
MATmin
MATmax
MMTmax
Seasonality
MATR
MMTR
MDTR
Isothermality
AB
DC
QR
PF
EC
ME
M
J
Y
B
V
P
S
SJ
W
PC2 ----------> Temperature Mean
27.20%
PC1 <---------- Temperature Variability
60.80%
% Variance Explained
2.5
5.0
7.5
10.0
B
Isolation Temperature
PC2
PC1
CTmax
CTmin
rmax
Topt
-7.5
-5.0
-2.5
0
2.5
5.0
t-value

**Supplementary Figure 15: Principal Components of environmental variation explain a small amount of Rate thermal trait variance. (A)** We obtained soil bioclimatic variables across sampling sites and ran a Principal Component Analysis (PCA) to compress their variability into orthogonal components. The first two PCs explained a cumulative 88.00% variance across sites. PC1 had a high negative correlation with metrics of temperature variability across temporal scales. PC2 was positively correlated with increases in mean annual temperatures, as well as minima and maxima. This allowed us to broadly categorize our sites based on their position relative to 0 along the range (PC1: Stable or Variable) and average (PC2: Cold or Hot) components, indicated by the text within the bigger circles. Each site abbreviation is connected to its coordinates on the surface of California. Site abbreviations by temperature profile are (Cold Variable): W - White Mountain Research Center, SJ - James San Jacinto Mountains Reserve, S - Sierra Nevada Aquatic Research Laboratory; (Cold Stable): V - Valentine Camp, Y - Yosemite Field Station, P - Point Reyes Field Station, B - Bodega Bay Research Center; (Hot Variable): AB - Steele/Burnand Anza-Borrego Desert Research Center Research Center, DC - Boyd Deep Canyon Desert Research Center, EC - Elliot Chaparral Reserve, QR - Quail Ridge Reserve, PF - Pinyon Flats; (Hot Stable): ME - Merced Vernal Pools and Grasslands Reserve, M - James McLaughlin Natural Reserve, J - Jepson Prairie Reserve. **(B)** We ran a hierarchical linear model that included the nested random effects of taxonomic groupings (Order and Genus), the two main principal components (PC1 and PC2), and the experimental isolation temperature as fixed effects, for each TPC trait. We extracted the t-value (Estimate / Standard Error) and partial R-squared (% Explained variance) for each variable. The t-value sign represents the sign of the association between a variable and a thermal trait, while the point size represents the trait variance explained by that variable. Overall, we see that experimental conditions explained a consistently higher fraction of trait variance than broad environmental variation components, and that the sign of the relationship depended on the variable and trait.

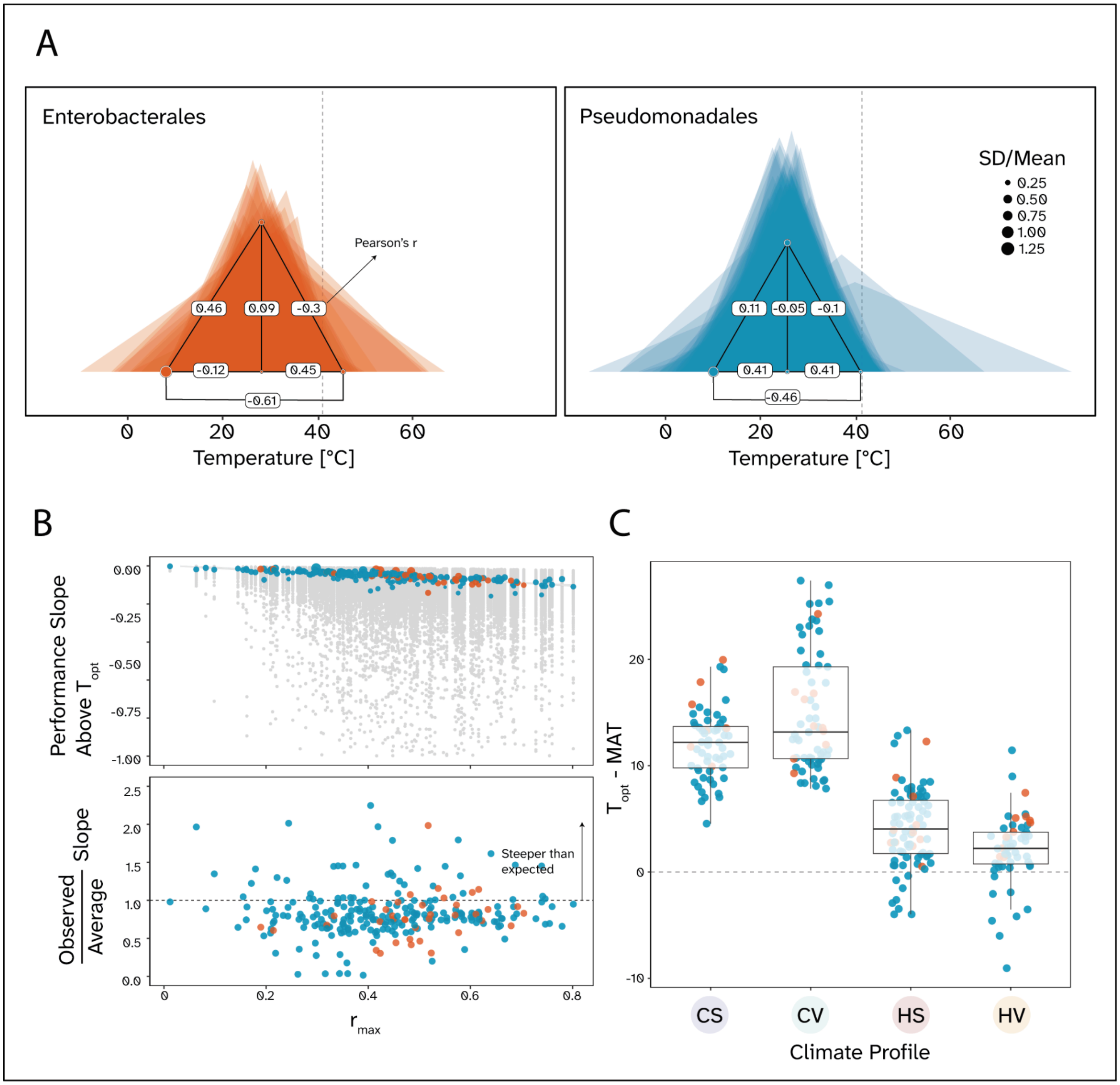

**Supplementary Figure 16: Variation in Growth Rate thermal performance shape is limited to changes in the lower thermal limit and peak performance. (A)** We generated simplified thermal performance curves for Enterobacterales (Orange) and Pseudomonadales (Blue), connecting the main traits represented by points (CTmin, CTmax, Topt, and Peak) with straight lines to form a triangle for each curve. The dashed line indicates the upper thermal limit documented for mesophilic enzymes at 42°C. Translucent triangles represent individual isolates, and full triangles represent average values by taxon. Point size is proportional to the trait variability within that taxon, as in SD(trait)/Mean(trait). The labels on the lines connecting traits represent the correlation (Pearson's r) coefficient between them. **(B)** We then calculated the slope of performance decrease above the thermal optimum of each simplified TPC. Here we show that the observed slope (colored points) increases (more negative = steeper) with performance peaks, suggesting the apparent robustness of the upper thermal limit (top). Compared to slopes obtained from a randomized distribution of thermal traits keeping the original peak values (grey points), the realized slopes land closer to the minimum possible values along the distribution, as represented by the ratio between the observed and average simulated slopes at each value of peak performance (bottom). **(C)** Finally, we evaluated the hypothesis that Jensen's inequality-based increases in Topt occur with temperature variability in cold- and hot-climate profiles. We observe contrasting trends: the difference between Topt and mean annual temperatures (MAT) increases with variability in generally cold environments, while it decreases in hot environments.

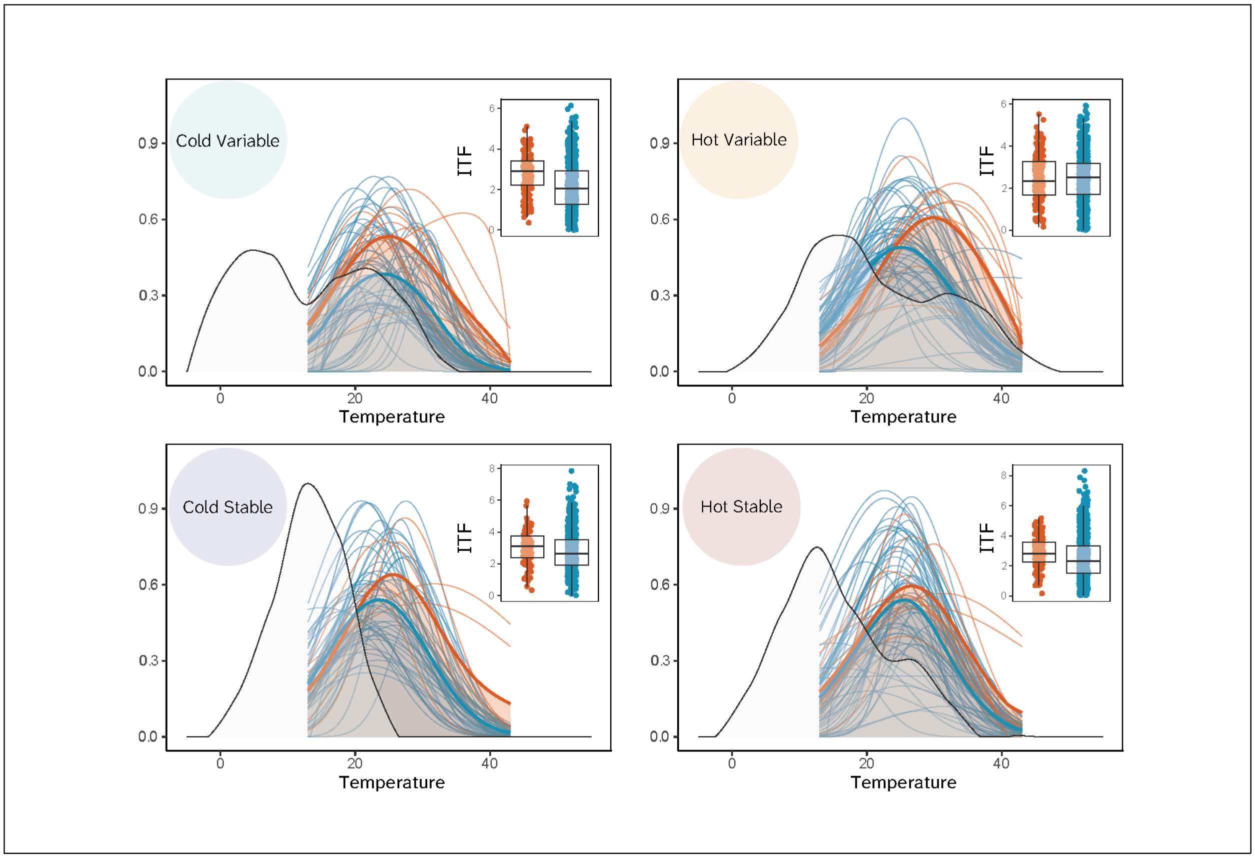

**Supplementary Figure 17: Overlaying the temperature probability by climate profile and the scaled performance of Rate by taxon shows differences in thermal fitness.** We obtained a probability distribution of temperatures (light grey) by climate profile during the plant growth season and overlaid a scaled performance of each taxon by temperature. Fine lines represent individual curves, and thick lines display average values for Enterobacterales (Orange) and Pseudomonadales (Blue). We then calculated the relative "Integrated Fitness" (inset) as the intersection area of each isolate curve with the temperature probability distribution.

**Supplementary Table 1: Overview of the bioclimatic variables per site obtained from publicly available databases.**

| Soil bioclimatic variable | Name and Abbreviation | Calculation | Units |
|---|---|---|---|
| SBIO1 | Annual mean temperature (MAT) | Mean (monthly ave) | °C |
| SBIO2 | Mean diurnal range (MMTR) | Mean (monthly max - monthly min) | °C |
| SBIO3 | Isothermality | (SBIO2/SBIO7) (×100) | |
| SBIO4 | Temperature seasonality | sd(monthly ave) | |
| SBIO5 | Maximum temperature of warmest month (MATmax) | max(monthly max) | °C |
| SBIO6 | Minimum temperature of coldest month (MATmin) | min(monthly min) | °C |
| SBIO7 | Temperature annual range (MATR) | (SBIO5-SBIO6) | °C |
| BIO12 | Annual Precipitation (Annual_precipitation) | Mean (annual ave) | mm |
| BIO13 | Precipitation of the wettest month (MAPmax) | max(monthly max) | mm |
| BIO14 | Precipitation of the driest month (MAPmin) | min(monthly min) | mm |
| BIO15 | Precipitation Seasonality (Precipitation_seasonality) | sd(monthly ave) | |

Supplementary Table 2: **Summary statistics of linear mixed regression models for each thermal trait**

| Trait | metric | Variable | Effect | Estimate | Std_Error | t_value | p_value | Partial _R2 | Variance_ob served | Lower_ CI | Upper _CI | Significant | Model |
|---|---|---|---|---|---|---|---|---|---|---|---|---|---|
| ctmax | auc | PC1 | fixed | 0 | 0.123 | -0.001 | 1 | 0 | - | - | - | - | lmerTest::lmer(ctmax ~ PC1 + PC2 + temperature_of_isolation + (1\|Order/Genus)) |
| ctmax | auc | PC2 | fixed | 0.674 | 0.168 | 4.006 | **0** | **0.051** | - | - | - | - | lmerTest::lmer(ctmax ~ PC1 + PC2 + temperature_of_isolation + (1\|Order/Genus)) |
| ctmax | auc | temperature_ of_isolation | fixed | 0.112 | 0.042 | 2.69 | **0.008** | **0.023** | - | - | - | - | lmerTest::lmer(ctmax ~ PC1 + PC2 + temperature_of_isolation + (1\|Order/Genus)) |
| ctmax | auc | Genus | random | - | - | - | - | - | 0.04 | 0 | 0.183 | FALSE | lmerTest::lmer(ctmax ~ PC1 + PC2 + temperature_of_isolation + (1\|Order/Genus)) |
| ctmax | auc | Order | random | - | - | - | - | - | **0.1** | **0** | **0.066** | **TRUE** | lmerTest::lmer(ctmax ~ PC1 + PC2 + temperature_of_isolation + (1\|Order/Genus)) |
| ctmax | auc | Residual | random | - | - | - | - | - | 0.86 | 0.782 | 1 | FALSE | lmerTest::lmer(ctmax ~ PC1 + PC2 + temperature_of_isolation + (1\|Order/Genus)) |
| ctmin | auc | PC1 | fixed | **-0.47** | **0.181** | **-2.595** | **0.01** | **0.016** | - | - | - | - | lmerTest::lmer(ctmin ~ PC1 + PC2 + temperature_of_isolation + (1\|Order/Genus)) |
| ctmin | auc | PC2 | fixed | -0.106 | 0.248 | -0.43 | 0.668 | 0.001 | - | - | - | - | lmerTest::lmer(ctmin ~ PC1 + PC2 + temperature_of_isolation + (1\|Order/Genus)) |
| ctmin | auc | temperature_ of_isolation | fixed | **-0.369** | **0.061** | **-6.023** | **0** | **0.109** | - | - | - | - | lmerTest::lmer(ctmin ~ PC1 + PC2 + temperature_of_isolation + (1\|Order/Genus)) |
| ctmin | auc | Genus | random | - | - | - | - | - | 0.062 | 0 | 0.127 | FALSE | lmerTest::lmer(ctmin ~ PC1 + PC2 + temperature_of_isolation + (1\|Order/Genus)) |
| ctmin | auc | Order | random | - | - | - | - | - | 0 | 0 | 0.074 | FALSE | lmerTest::lmer(ctmin ~ PC1 + PC2 + temperature_of_isolation + (1\|Order/Genus)) |
| ctmin | auc | Residual | random | - | - | - | - | - | 0.938 | 0.873 | 1 | FALSE | lmerTest::lmer(ctmin ~ PC1 + PC2 + temperature_of_isolation + (1\|Order/Genus)) |
| rmax | auc | PC1 | fixed | **0.048** | **0.023** | **2.061** | **0.04** | **0.004** | - | - | - | - | lmerTest::lmer(rmax ~ PC1 + PC2 + temperature_of_isolation + (1\|Order/Genus)) |

| | | | | | | | | | | | | | |
|---|---|---|---|---|---|---|---|---|---|---|---|---|---|
| rmax | auc | PC2 | fixed | 0.012 | 0.032 | 0.392 | 0.695 | 0 | - | - | - | - | lmerTest::lmer(rmax ~ PC1 + PC2 + temperature_of_isolation + (1\|Order/Genus)) |
| rmax | auc | temperature_of_isolation | fixed | **-0.021** | **0.008** | **-2.719** | **0.007** | **0.006** | - | - | - | - | lmerTest::lmer(rmax ~ PC1 + PC2 + temperature_of_isolation + (1\|Order/Genus)) |
| rmax | auc | Genus | random | - | - | - | - | - | **0.321** | **0** | **0.129** | **TRUE** | lmerTest::lmer(rmax ~ PC1 + PC2 + temperature_of_isolation + (1\|Order/Genus)) |
| rmax | auc | Order | random | - | - | - | - | - | **0.485** | **0** | **0.059** | **TRUE** | lmerTest::lmer(rmax ~ PC1 + PC2 + temperature_of_isolation + (1\|Order/Genus)) |
| rmax | auc | Residual | random | - | - | - | - | - | **0.194** | **0.845** | **1** | **TRUE** | lmerTest::lmer(rmax ~ PC1 + PC2 + temperature_of_isolation + (1\|Order/Genus)) |
| topt | auc | PC1 | fixed | **-0.215** | **0.091** | **-2.353** | **0.019** | **0.01** | - | - | - | - | lmerTest::lmer(topt~ PC1 + PC2 + temperature_of_isolation + (1\|Order/Genus)) |
| topt | auc | PC2 | fixed | **0.424** | **0.123** | **3.44** | **0.001** | **0.022** | - | - | - | - | lmerTest::lmer(topt~ PC1 + PC2 + temperature_of_isolation + (1\|Order/Genus)) |
| topt | auc | temperature_of_isolation | fixed | **-0.142** | **0.03** | **-4.73** | **0** | **0.041** | - | - | - | - | lmerTest::lmer(topt~ PC1 + PC2 + temperature_of_isolation + (1\|Order/Genus)) |
| topt | auc | Genus | random | - | - | - | - | - | **0.465** | **0** | **0.168** | **TRUE** | lmerTest::lmer(topt~ PC1 + PC2 + temperature_of_isolation + (1\|Order/Genus)) |
| topt | auc | Order | random | - | - | - | - | - | 0 | 0 | 0.051 | FALSE | lmerTest::lmer(topt~ PC1 + PC2 + temperature_of_isolation + (1\|Order/Genus)) |
| topt | auc | Residual | random | - | - | - | - | - | **0.535** | **0.822** | **1** | **TRUE** | lmerTest::lmer(topt~ PC1 + PC2 + temperature_of_isolation + (1\|Order/Genus)) |
